# Distinct roles for CDK-Mediator in controlling Polycomb-dependent chromosomal interactions and priming genes for induction

**DOI:** 10.1101/2021.11.04.467119

**Authors:** Emilia Dimitrova, Angelika Feldmann, Robin H. van der Weide, Koen D. Flach, Anna Lastuvkova, Elzo de Wit, Robert J. Klose

## Abstract

Precise control of gene expression underpins normal development. This relies on mechanisms that enable communication between gene promoters and other regulatory elements. In embryonic stem cells (ESCs), the CDK-Mediator (CDK-MED) complex has been reported to physically link gene regulatory elements to enable gene expression and also prime genes for induction during differentiation. Here we discover that CDK-MED contributes little to 3D genome organisation in ESCs, but has a specific and essential role in controlling interactions between inactive gene regulatory elements bound by Polycomb repressive complexes (PRCs). These interactions are established by the canonical PRC1 (cPRC1) complex but rely on CDK-MED, which facilitates binding of cPRC1 to its target sites. Importantly, through separation of function experiments, we reveal that this collaboration between CDK-MED and cPRC1 in creating long-range interactions does not function to prime genes for induction during differentiation. Instead, we discover that priming relies on an interaction-independent mechanism whereby the CDK module supports core Mediator engagement with gene promoters to enable gene activation.

## INTRODUCTION

Mechanisms that shape 3D genome organisation are thought to play important roles in controlling gene expression, particularly during development. For example, interactions between gene promoters, or gene promoters and other distal gene regulatory elements, like enhancers, have been implicated both in maintenance of gene expression patterns and in enabling alterations in gene expression states during cell fate transitions ^1-4^.

A number of mechanisms have been proposed to create and regulate interactions between gene regulatory elements. For example, cohesin can extrude chromatin to establish topologically associated domains (TADs), which are generally constricted by insulator sites bound by CTCF. Cohesin-mediated loop extrusion is thought to increase the frequency of interactions between gene promoters and distal regulatory elements within TADs ^5^. However, while the disruption of cohesin or CTCF has profound effects on interactions within TADs, this typically translates into modest or tissue-specific effects on gene expression ^6-10^. While loop extrusion functions across the genome, other mechanisms are also thought to play more direct and specific roles at gene regulatory elements by creating physical interactions that may control gene expression. For example, the Mediator complex, which is a fundamental regulator of gene transcription, has been proposed to support gene expression by functioning as a molecular bridge through binding transcription factors at active enhancers and RNA Polymerase II at gene promoters ^11-13^. However, recent work has questioned the extent to which the function of Mediator in gene expression relies on promoting physical interactions between regulatory elements ^14-17^. At silent gene regulatory elements, binding of the Polycomb repressive complexes (PRCs) enables physical interactions between these inactive sites ^18-28^, which is thought to maintain gene repression ^29-31^ but may also poise genes for activation during cell linage commitment ^32-34^. In these contexts, whether chromosomal interactions themselves or other functions of the Polycomb system control gene expression is unknown. Therefore, although it is clear that a variety of mechanisms have evolved to shape how gene regulatory elements physically interact with one another, the extent to which this is required to control gene expression remains a central outstanding question in the field ^35-40^.

Although it appears that the Mediator complex alone may not play a major role in enabling interactions between gene regulatory elements ^14-17^, we and others have shown that a distinct form of the complex, containing the cyclin-dependent kinase module (CDK) module (composed of CDK8/19, CNCC, MED12/12L, and MED13/13L) which does not interact with RNA polymerase II, is associated with gene regulatory element interactions in mouse embryonic stem cells (ESCs) ^41-46^. Unlike Mediator, CDK-MED has been implicated in both repressing and supporting gene expression, suggesting that it might work through mechanisms that are distinct from the well-characterised function of Mediator in binding to and regulating RNA Polymerase II activity ^47,48^. In line with this possibility, CDK-MED appears to play specialised roles in controlling inducible gene expression after exposure to extracellular stimuli or cellular differentiation cues ^47,49-54^. We and others have previously demonstrated that CDK-MED is recruited to the promoters of repressed developmental genes in ESCs ^55-58^ and this primes these genes for induction during differentiation ^55^. In this context, CDK-MED binding appears to be important for creating interactions with other gene regulatory elements, suggesting that formation of 3D interactions may underpin its capacity to prime developmental genes for induction during cell lineage commitment ^41^.

Based on these findings, we set out to determine how CDK-MED controls chromosomal interactions and gene expression. To achieve this, we exploit inducible genetic perturbation systems and genomic approaches to examine CDK-MED function in ESCs and during cellular differentiation. We discover that CDK-MED contributes little to overall 3D genome organisation in ESCs, but is essential for creating interactions between Polycomb-bound regions of the genome. We show that CDK-MED does not define these interactions through an intrinsic bridging mechanism. Instead, it controls canonical Polycomb repressive complex 1 (cPRC1) binding at these sites, which in turn establishes contacts between Polycomb domains. Surprisingly, through separation of function experiments we reveal that Polycomb-dependent chromosomal interactions regulated by CDK-MED are not required for the priming/poising of genes for induction during differentiation. Instead, we discover that the priming function of CDK-MED relies on its ability to enable core Mediator binding to gene promoters during the process of gene induction.

## RESULTS

### CDK-MED has a limited role in 3D genome organisation but is essential for Polycomb domain interactions

To examine how CDK-MED influences genome organisation in ESCs, we carried out *in situ* Hi-C in a cell line where we can inducibly disrupt CDK-MED complex formation by removing its MED13/MED13L structural subunits (CDK-MED cKO, Figure 1A-B, S1A) ^55^. Importantly we observed no major alterations to overall genome organisation after CDK-MED disruption, with TADs and loop interactions remaining largely unchanged (Figure 1C, D). Previously it has been proposed that CDK-MED could promote super enhancer-promoter interactions in ESCs ^42,43^. However, we observed only subtle reductions in these interactions upon disruption of CDK-MED (Figure S1C). Therefore, we conclude that CDK-MED does not contribute centrally to 3D genome organisation in ESCs.

**Figure 1:**
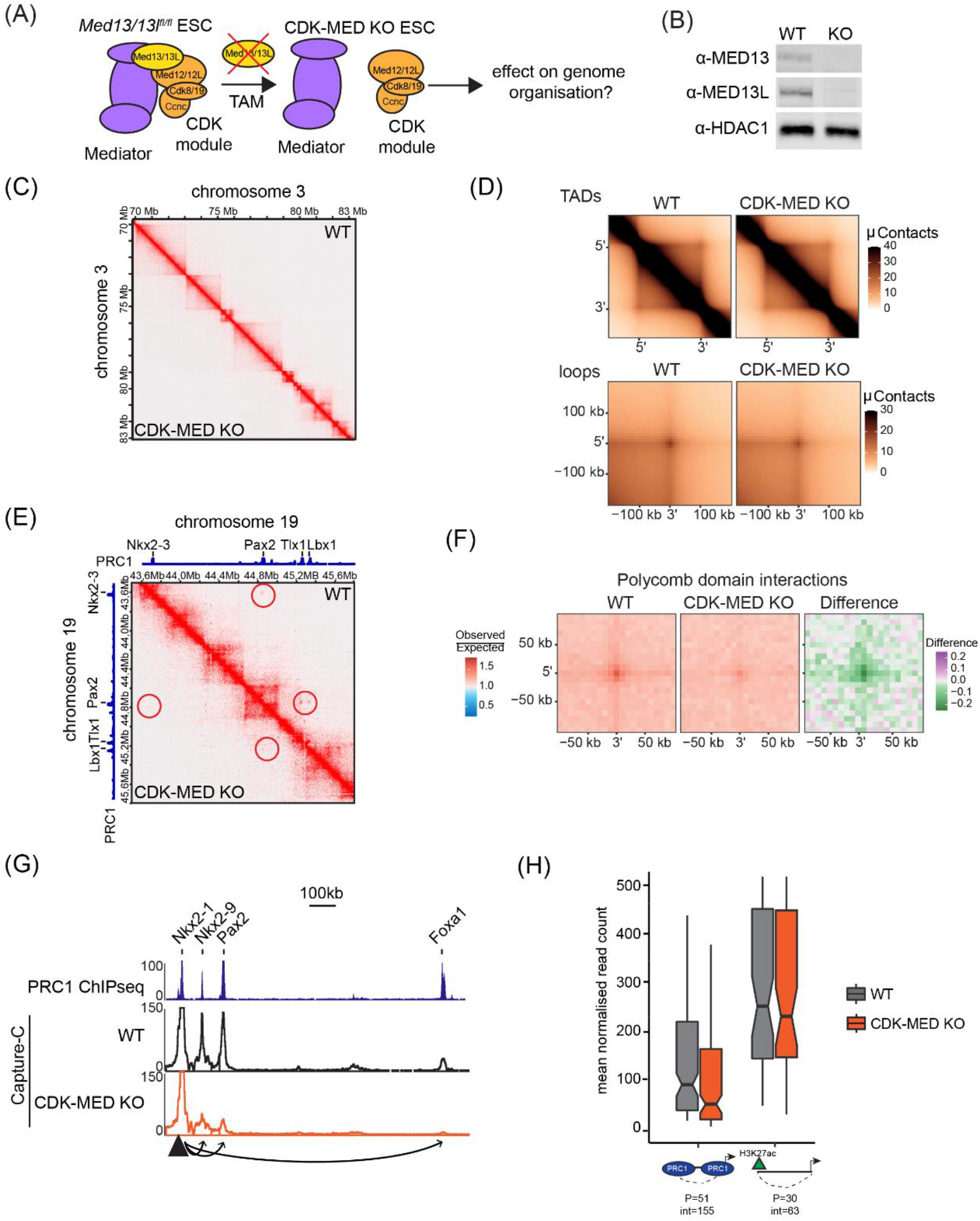
CDK-MED has a limited role in 3D genome organisation but is essential for Polycomb domain interactions. (A) A schematic illustration of *Med13/13l*^*fl/fl*^ ESCs where 4-hydroxytamoxifen (TAM) induces conditional disruption of the CDK-MED complex. (B) Western blot analysis of nuclear extracts from *Med13/13l*^*fl/fl*^ (WT) and *Med13/13l*^*-/-*^ (CDK-MED KO) ESCs showing deletion of MED13 and MED13L proteins. HDAC1 is shown as a loading control. (C) Hi-C contact matrices of WT and CDK-MED KO ESCs at 10kb resolution. Genomic coordinates are indicated. (D) Aggregate analysis of TADs (top) and loops (bottom) in WT and CDK-MED KO ESCs at 100kb resolution. (E) Hi-C contact matrices of WT and CDK-MED KO ESCs at 5kb resolution. Interactions between Polycomb domains are indicated with a red circle. The blue track shows binding of PRC1 (RING1B ChIPseq). Genomic coordinates are indicated. (F) Aggregate analysis of interactions between PRC1 peaks (RING1B) within Polycomb domains in WT and CDK-MED KO ESCs at 50kb resolution. The difference between WT and KO is shown. (G) A snapshot showing Capture-C read count signal in WT and CDK-MED KO ESCs. Interactions between the *Nkx2-1* promoter bait (triangle) and surrounding Polycomb-bound sites are shown with arrowheads. PRC1 binding (RING1B ChIPseq) is shown as a reference. (H) Boxplot analysis of mean normalised read count from WT and CDK-MED KO ESCs showing interactions between Polycomb gene promoters with other Polycomb-domains (left) or non-Polycomb gene promoters with active sites (H3K27ac, right). Number of promoters (P) and interactions (int) is shown.

ESCs are characterised by a unique set of extremely strong long-range interactions between regions of the genome that have high level occupancy of Polycomb repressive complexes, which we refer to as Polycomb domains ^19-21,23,28,59^. These interactions are thought to contribute to developmental gene regulation either by maintaining repression in differentiated cell types or potentially by poising genes for induction during cell lineage commitment. Interestingly, a similar role in regulating developmental gene expression has also been proposed for CDK-MED ^41,55-57^. Given these seemingly similar functionalities, we asked whether CDK-MED might influence interactions between Polycomb domains. Remarkably, Hi-C analysis after CDK-MED disruption revealed dramatic reductions in interactions between Polycomb domains (Figure 1E-F). We also confirmed this effect using Capture-C analysis focussed on promoters associated with Polycomb domains (Figure 1G-H, S1D). Therefore, we discover that CDK-MED is essential for interactions between Polycomb domains.

### CDK-MED regulates canonical PRC1 binding to enable interactions between Polycomb domains

To understand how CDK-MED enables interactions between Polycomb domains, we asked if CDK-MED is bound at these sites. We found that the majority of Polycomb domains (91.12%) were also enriched for the CDK-MED subunit CDK8, in general agreement with previous findings ^56,57^, suggesting that its effects on Polycomb domain interactions may be direct (Figure 2A, S2A). It has previously been proposed that interactions between Polycomb domains are dependent on the canonical Polycomb Repressive Complex 1 (cPRC1), which is defined by its structural subunit PCGF2 ^20,24,30,60-65^. Given the profound effects on Polycomb domain interactions upon loss of CDK-MED, we hypothesised that CDK-MED may influence the function of cPRC1. To test this possibility, we examined cPRC1 occupancy after CDK-MED disruption by carrying out calibrated ChIP-seq (cChIP-seq) for its subunits RING1B and PCGF2. Importantly, this revealed a major reduction in cPRC1 binding at Polycomb target sites in the absence of CDK-MED (Figure 2B-D, S2B-C), despite only subtle reductions in protein levels (Figure S2D). cPRC1 associates with Polycomb domains via its CBX7 subunit that binds H3K27me3 deposited by PRC2 ^66-68^. Interestingly, cChIP-seq for H3K27me3 revealed only modest reductions in this modification after CDK-MED disruption (Figure 2B-D). Therefore, CDK-MED regulates cPRC1 binding without major effects on H3K27me3.

**Figure 2:**
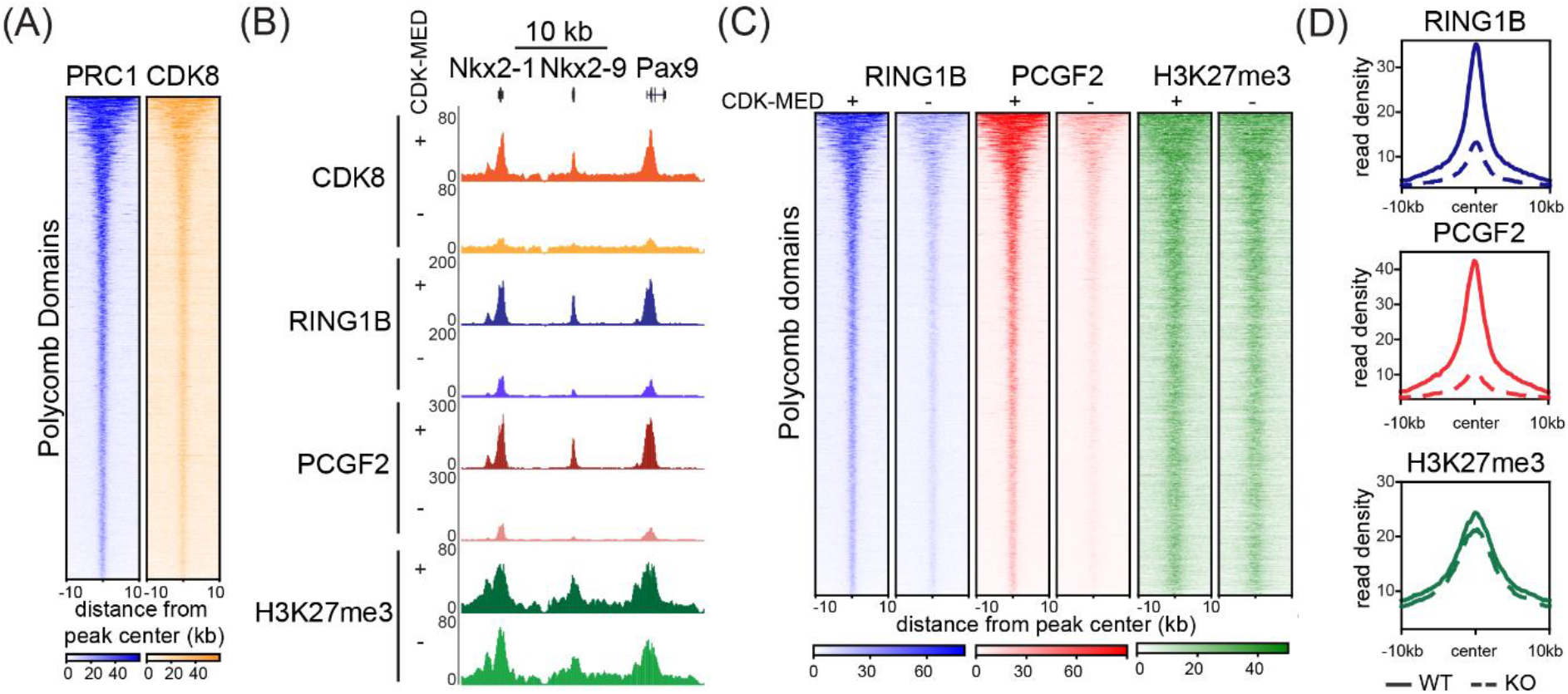
CDK-MED regulates canonical PRC1 binding. (A) Heatmaps showing RING1B (PRC1) and CDK8 ChIPseq signal at Polycomb domains (n=2097), sorted by decreasing RING1B signal. (B) A genomic snapshot of a Polycomb-bound locus, showing CDK8, RING1B, PCGF2 and H3K27me3 ChIPseq signal in WT (+) and CDK-MED KO (-) ESCs. (C) Heatmaps showing RING1B, PCGF2 and H3K27me3 ChIPseq signal at Polycomb domains (n=2097) in WT (+) and CDK-MED KO (-) ESCs, sorted by decreasing RING1B signal. (D) Metaplot analysis of RING1B, PCGF2 and H3K27me3 enrichment at Polycomb domains (n=2097) in WT and CDK-MED KO ESCs.

Given that cPRC1 has been proposed to enable interactions between Polycomb domains ^20,24,30^, and its binding is abrogated following disruption of CDK-MED (Figure 2), the observed effect on Polycomb domain interactions in the absence of CDK-MED may be due to loss of cPRC1 occupancy. However, CDK-MED has also been proposed to function as a molecular bridge to enable chromosomal interactions ^41-43^. Given that both cPRC1 and CDK-MED binding will be lost upon CDK-MED disruption, interactions could be defined by either cPRC1 or CDK-MED. To discover the molecular determinant that enables these interactions, we took advantage of a synthetic system to create a separation of function scenario where either cPRC1 or CDK-MED could be ectopically tethered to an artificial site in the genome ^69^ (Figure 3A, S3A-B). Tethering of PCGF2 recruits the cPRC1 complex ^69^ and tethering of CDK8 recruits CDK-MED (Figure S3C). We then asked whether binding of cPRC1 or CDK-MED at this ectopic site was able to support interactions with nearby regions co-occupied by cPRC1 and CDK-MED. This revealed that cPRC1 was sufficient to create *de novo* interactions with surrounding sites in line with similar findings from PRC2 tethering ^70^, which would lead to recruitment of cPRC1 ^69^. In contrast, we found no evidence for interactions with surrounding sites when CDK-MED was tethered (Figure 3B, S3D). Importantly, endogenous control sites retained interactions in both cell lines, although they were slightly weaker in the CDK-MED tethered line (Figure S3E).

**Figure 3:**
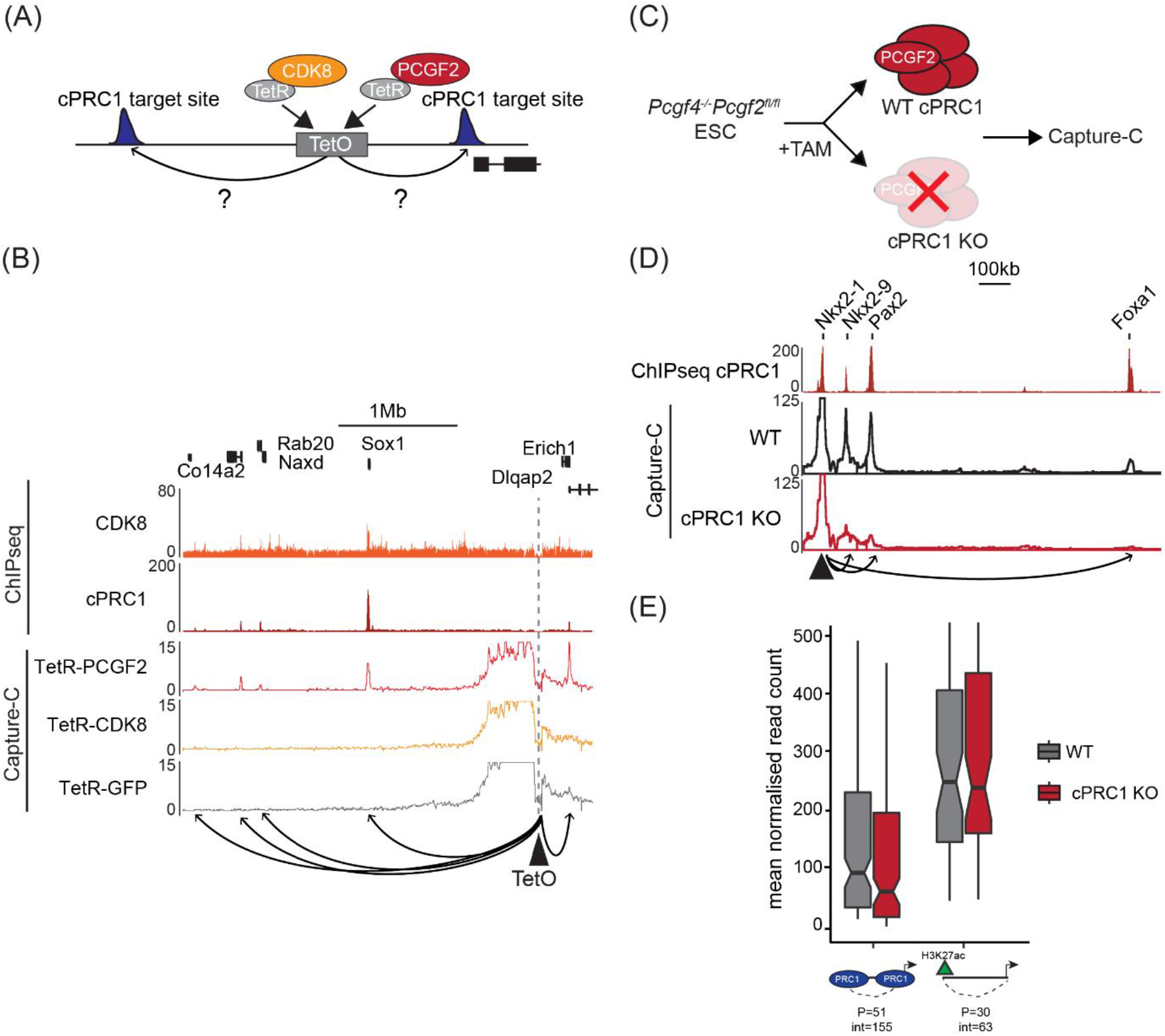
cPRC1 creates interactions between Polycomb domains. (A) A schematic illustration of the integrated TetO site and experimental setup. (B) A snapshot showing Capture-C read count signal from TetR-PCGF2, TetR-CDK8 and TetR-GFP lines at the TetO array. CDK8 and PCGF2 (cPRC1) ChIPseq signal is given as a reference. The TetO bait is shown as a triangle and interactions created with surrounding cPRC1-bound sites are represented with arrowheads. (C) A schematic illustration of the cPRC1 (*Pcgf4*^*-/-*^*Pcgf2*^*fl/fl*^) conditional knock-out line. (D) A snapshot showing Capture-C read count signal from WT and cPRC1 KO ESCs. Interactions between the *Nkx2-1* promoter bait (triangle) and surrounding Polycomb domain sites are shown with arrowheads. cPRC1 binding (PCGF2 ChIPseq) is shown as a reference. (E) Boxplot analysis of normalised read counts from WT and cPRC1 KO ESCs showing interactions between Polycomb gene promoters with other Polycomb-domains (left) or non-Polycomb gene promoters with active sites (H3K27ac, right). Number of promoters (P) and interactions (int) is shown.

To further explore whether cPRC1 is the central determinant underpinning Polycomb domain interactions, we next used a cell line in which we can inducibly disrupt the cPRC1 complex by removing the core structural components PCGF2/4 (cPRC1 cKO) ^59^ and carried out Capture-C (Figure 3C, S3F). Importantly, removal of cPRC1 caused a near complete loss of interactions between Polycomb domains, while most sites retained CDK-MED binding (Figure 3D-E, S3G-I). Therefore, cPRC1 establishes long-range interactions between Polycomb domains, with CDK-MED playing a regulatory role in facilitating cPRC1 binding.

### CDK-MED primes genes for activation during differentiation independently of cPRC1-mediated interactions

CDK-MED occupies silent developmental gene promoters in ESCs and this is required for their subsequent activation during differentiation ^55,56^. In some cases this corresponded to pre-formed long-range interactions with other gene regulatory elements, suggesting that by bringing gene regulatory elements into close proximity in ESCs, CDK-MED could prime these for future activation ^41^. We now discover that CDK-MED-dependent interactions rely on cPRC1 (Figure 3). Importantly, the Polycomb system has similarly been implicated in poising or priming genes for activation during differentiation via creating interactions between gene promoters and other regulatory elements, including poised enhancers ^32,33^. Based on this functional convergence of CDK-MED and cPRC1 activities, we hypothesised that CDK-MED may enable the formation of interactions between gene promoters and regulatory elements via a cPRC1-dependent mechanism to prime genes for activation during differentiation.

To examine this possibility, we used all-trans retinoic acid (RA) to drive ESC differentiation and carried out calibrated nuclear RNA-seq (cnRNA-seq) to identify genes that rely on CDK-MED for their induction during differentiation (Figure 4A, S4A-B). Based on this analysis, we identified 631 (fold change >1.5, padj<0.05) CDK-MED-dependent genes (Figure 4B, S4C). Importantly these genes also showed enrichment for cPRC1 at their promoters in ESCs (Figure S4D). To determine whether cPRC1 and its capacity to mediate chromosomal interactions would enable gene induction by CDK-MED, we depleted cPRC1 and induced differentiation (Figure 4C, S4E-F). Importantly, this revealed that on average CDK-MED-dependent genes induced normally in the absence of cPRC1 (Figure 4D-E, S4G), with only 18 of these genes showing a significant decrease in activation (Figure S4G-H). Therefore, while CDK-MED contributes to gene induction, we find no evidence that it does so via a cPRC1-dependent mechanism.

**Figure 4:**
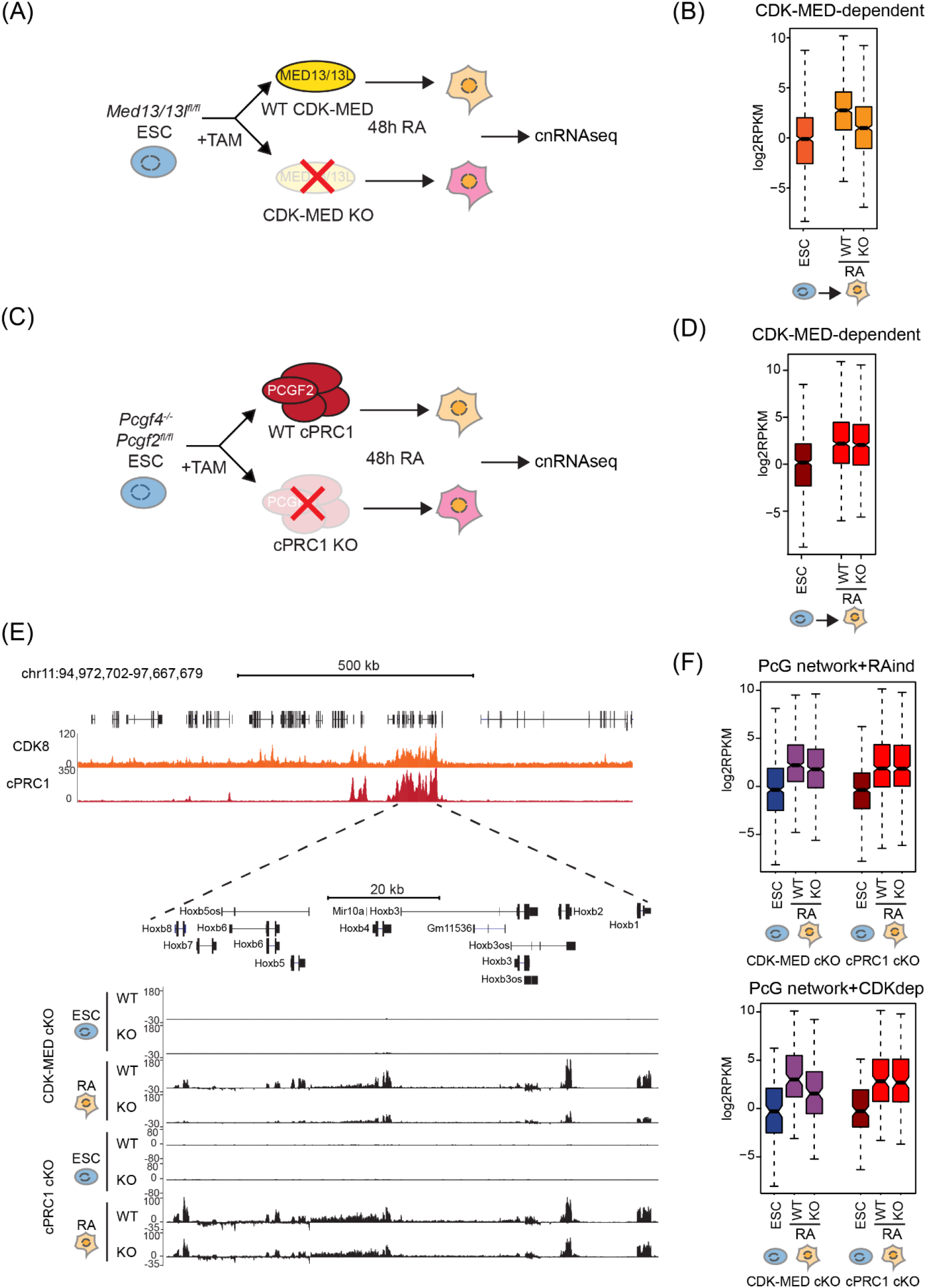
CDK-MED primes genes for activation during differentiation independently of cPRC1-mediated interactions. (A) A schematic illustration of the differentiation of WT and CDK-MED KO ESCs for cnRNAseq. (B) Boxplot analysis of the expression of CDK-MED-dependent genes (n=631) in WT ESCs and following RA-induction (WT and CDK-MED KO). (C) A schematic illustration of the differentiation of WT and cPRC1 KO ESCs for cnRNAseq; (D) As in (B) for cPRC1 cKO cells. (E) A screen-shot showing the expression of genes within the *HoxB* cluster following RA induction of CDK-MED cKO or cPRC1 KO cells. Forward strand is shown on top and reverse strand is shown at the bottom of each track. ChIPseq tracks for CDK8 and cPRC1 (PCGF2) enrichment are shown. (F) Boxplot analysis of the expression of RA-induced genes from the Polycomb (PcG) network (top) and CDK-MED-dependent genes from the PcG network (bottom) following RA induction of CDK-MED cKO or cPRC1 KO cells.

This finding prompted us to analyse more generally whether cPRC1 has a role in gene induction during differentiation, particularly of genes that engage in interactions. To achieve this, we extended our analysis to RA-induced genes that are part of a previously described Polycomb interaction network in ESCs (n=482) (Figure S4I-J) ^19^. Importantly, we confirmed that interactions between these genes are lost in the absence of cPRC1, including interactions with poised enhancers (Figure S4K-L). However, as with CDK-MED-dependent genes, this had minimal effect on gene induction (Figure 4E-F, S4M). In contrast, we identified 184 genes within the Polycomb interaction network that rely on CDK-Mediator for induction (Figure 4F). Therefore, we discover that CDK-MED has an essential role in gene activation during differentiation, but this is independent of cPRC1-mediated chromosomal interactions. Furthermore, we show that cPRC1 does not poise genes for activation during differentiation, despite its role in enabling interactions between gene promoters and other regulatory elements in ESCs.

### CDK-MED primes genes for activation via recruitment of the core Mediator complex

Although CDK-MED is essential for enabling cPRC1 to create interactions between Polycomb domain-associated gene regulatory elements, these interactions are dispensable for gene induction during differentiation. In the absence of a pre-formed interaction mechanism for priming, we hypothesised that CDK-MED may prime genes for activation during differentiation by more directly controlling the function of the core Mediator ^71-73^. To investigate this possibility, we engineered an epitope tag into the endogenous *Med14* gene, which is a structural subunit of the core Mediator, and carried out ChIP-seq analysis to examine its occupancy in ESCs and during differentiation (Figure S5A-C). Interestingly, unlike CDK8, MED14 was depleted from the promoters of CDK-dependent genes (Figure 5) and, more broadly, from Polycomb domains in ESCs (Figure S5D). This suggests that the CDK module can bind to inactive developmental gene promoters independently of stable binding with core Mediator as has been suggested previously ^58^.

**Figure 5:**
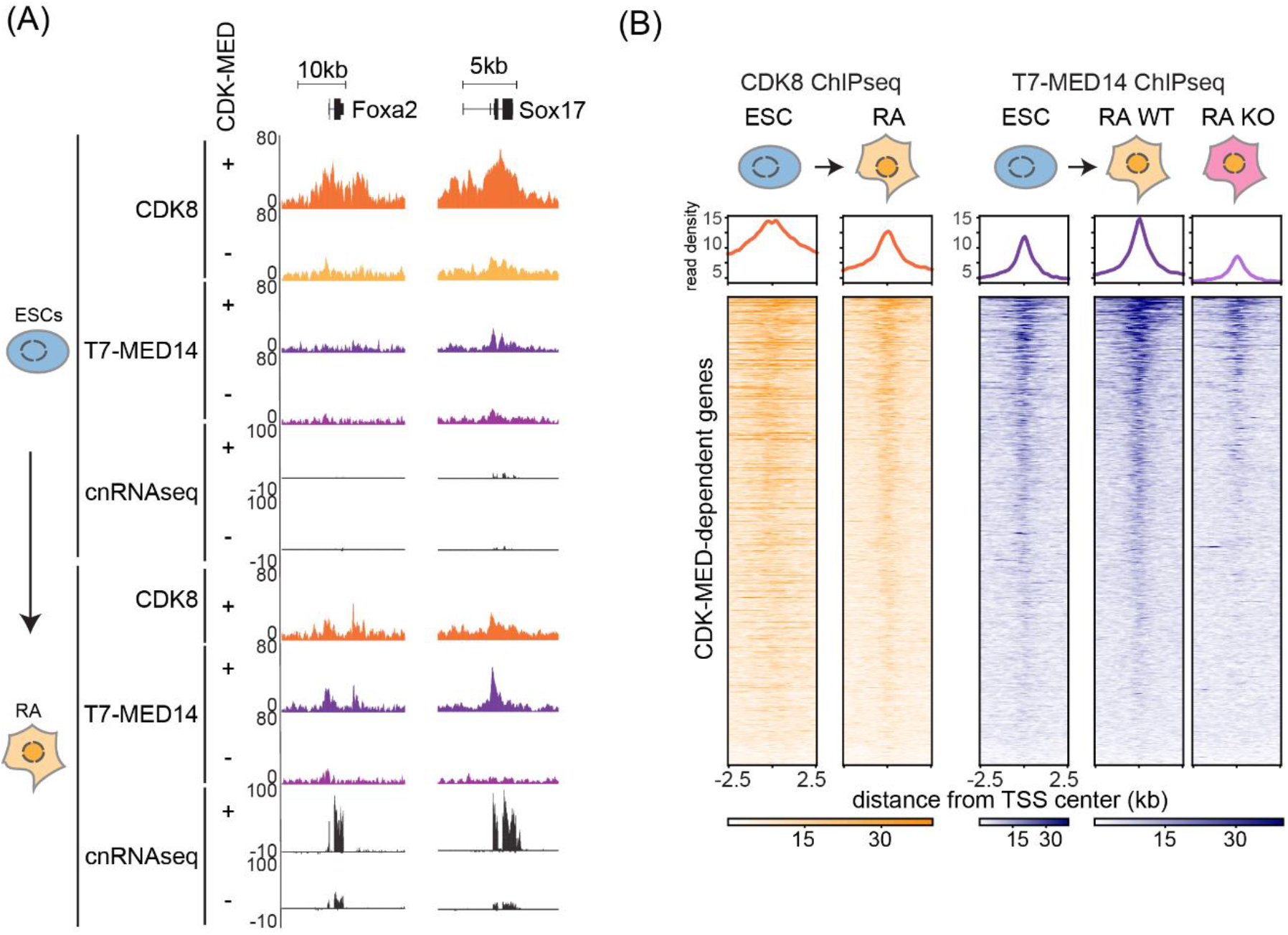
CDK-MED enables gene induction via recruitment of the Mediator complex. (A) A genomic snapshot of two CDK-MED-dependent genes, showing CDK8 and T7-MED14 ChIPseq and cnRNAseq in WT (+) and CDK-MED KO (-) ESCs (top) and following RA-induction (bottom). (B) Heatmaps showing CDK8 and T7-MED14 ChIPseq signal at promoters (+/- 2.5kb) of CDK-MED-dependent genes in ESCs and following RA-induction (n=631). T7-MED14 signal is shown for WT and CDK-MED KO RA-induced cells. Genes are sorted by decreasing T7-MED14 signal in RA-treated cells. Metaplots showing read density are shown on top of each heatmap.

Based on these observations, we were keen to examine core Mediator association with these sites during differentiation. During the transition to an active state, promoters of CDK-MED-dependent genes showed reduced levels of CDK8 binding and acquired MED14 (Figure 5, S5E-F). We then asked whether the CDK module was required for the transition from a CDK-predominating to a core Mediator-predominating state during differentiation ^12,13,58^. This revealed that following differentiation, the promoters of CDK-MED-dependent genes do not acquire MED14 in the absence of the CDK module (Figure 5A-B, S5F), consistent with these genes failing to induce appropriately (Figure 4). Therefore, we propose that the CDK module primes genes for induction not by pre-forming 3D gene regulatory interactions through the Polycomb system, but instead by enabling efficient engagement of the core Mediator at target gene promoters to drive transcription activation during differentiation.

## DISCUSSION

Defining the extent to which interactions between gene regulatory elements are required for controlling gene expression has been challenging. This is due to the fact that many of the proteins and complexes that are proposed to enable these interactions are also known to have direct roles in controlling transcription at gene promoters. Here we discover that CDK-MED contributes very little to 3D genome organisation in ESCs but is specifically required for interactions between Polycomb-bound gene regulatory elements (Figure 1). These interactions do not rely directly on a CDK-MED-based bridging mechanism (Figure 3), but instead CDK-MED controls binding of the cPRC1 complex (Figure 2), which enables interactions between Polycomb domains (Figure 3). By removing cPRC1, we specifically disrupt these interactions, yet reveal that CDK-MED is still able to prime genes for activation during differentiation (Figure 4), through supporting recruitment of the core Mediator to gene promoters (Figure 5). Therefore, CDK-MED primes genes for activation during differentiation through recruitment of the core Mediator.

Physical interactions between gene regulatory elements are thought to enable gene expression ^32,41,74,75^. In line with this concept, it was seductive to propose that, through the function of Polycomb and/or CDK-MED complexes, pre-formed interactions that tether silent developmental genes and other regulatory elements in stem cells may render genes poised or primed for activation during differentiation ^32-34,41,76,77^. Here we demonstrate that pre-formed interactions between gene regulatory elements co-occupied by CDK-MED and cPRC1 rely on cPRC1, and that the binding of cPRC1 is regulated by CDK-MED. While the precise mechanisms through which CDK-MED facilitates cPRC1 binding to create these interactions remains an open question for further studies, this realisation allowed us to create a separation of function scenario whereby we could disrupt pre-formed interactions by removing cPRC1, yet leave CDK-MED intact. Importantly, in the context of these experiments, we find no evidence to suggest that pre-formed regulatory interactions play a prominent role in priming genes for activation during differentiation. In line with these interactions having a limited role in gene activation, cPRC1 also does not contribute to gene regulation during embryoid body formation *in vitro* ^78^ and cPRC1-null mice develop normally until 8.5 dpc, when a host of key developmental gene expression transitions have already completed ^79,80^. Instead, we find that the CDK module appears to have a more direct role in priming genes for induction during differentiation and does so by ensuring appropriate binding of the core Mediator complex during gene activation. This priming likely involves the CDK-MED interaction partner FBXL19 that directly binds CpG islands and as such allows the CDK module to associate with silent developmental gene promoters ^55^. We envisage that pre-binding of the CDK module could provide a platform on which the core Mediator could dynamically interact and that other transcriptional activators may function in concert with the CDK module to stabilise core Mediator binding during gene induction. In line with this view, removal of FBXL19 causes a reduction in CDK module binding at silent developmental gene promoters and, similarly to CDK-MED removal, renders them less competent for induction during differentiation ^55^. Furthermore, mice deficient for CDK-module subunits display pre-implantation lethality, consistent with an essential role in early developmental gene expression transitions ^81-83^. As such, CDK-Mediator appears to function to prime genes for induction through regulating core Mediator function at gene promoters, not through mechanisms that create pre-formed regulatory element interactions.

These new findings then raise the important question of why CDK-MED regulates cPRC1 binding to create interactions between silent gene regulatory elements if this is not related to its role in priming genes for induction during differentiation. A hint as to why this might be important comes from genetic screens in *Drosophila* where the CDK-MED complex components MED12 and MED13 were identified as Polycomb group genes that enable the long-term maintenance of *Hox* gene repression ^84^. In agreement with a potential repressive role for CDK-MED at Polycomb target genes, it was recently shown that the CDK8 component of the CDK-MED complex has important roles in maintaining X-chromosome inactivation in mice ^85^, and that the absence of CDK8 led to loss of Polycomb-mediated gene silencing ^85,86^. Interestingly, in both of these scenarios, CDK-MED and Polycomb appear to maintain repression in more differentiated cells, while, in contrast, cPRC1 disruption has little effect on the maintenance of Polycomb target gene repression in ESCs ^59,78,87^. As such, we envisage that the role CDK-MED plays in regulating cPRC1 occupancy to create long-range interactions between silent regulatory elements may be particularly important in maintaining long-term gene repression in more differentiated cell types and less important in rapidly dividing stem cells. This is also consistent with cPRC1-deficient mice displaying inappropriate maintenance of Polycomb target gene repression and lethality in later embryonic stages ^79,80^.

Based on its seemingly distinct roles in gene regulation, we propose that CDK-MED may play a ‘yin- and-yang’ role in controlling expression. We envisage that, during early developmental stages, the CDK-module associates with silent developmental gene promoters to support gene induction during differentiation by helping to stabilise core Mediator binding during the transition to an activated state. However, in the absence of an activation signal at later developmental stages, its distinct role in enabling cPRC1 binding to create interactions with other silent Polycomb-occupied regulatory sites could predominate in helping to maintain long-term gene repression. As such, these distinct CDK-MED activities could play important and complementary roles in supporting developmental gene regulation. In future work it will be important to test these new models for CDK-MED function in appropriate mouse developmental model systems.

In summary, we discover that CDK-MED is essential for regulating interactions between Polycomb domains. However, these interactions contribute very little to gene activation during differentiation. Instead, we show that CDK-MED primes genes for induction during differentiation through supporting core Mediator biding to promoters upon gene activation.

## MATERIALS AND METHODS

### Cell culture

Mouse ESCs were cultured on gelatine-coated (Sigma) dishes in Dulbecco’s Modified Eagle Medium (DMEM, Thermo Fisher scientific) supplemented with 15% fetal bovine serum (BioSera), 2mM L-Glutamine, 0.5mM beta-mercaptoethanol, 1x non-essential amino acids, 1x penicillin/streptomycin solution (Thermo Fisher scientific) and 10 ng/mL leukemia-inhibitory factor (produced in-house). *Med13/13l*^*fl/fl*^ ERT2-Cre ESCs ^55^ and *Pcgf4*^*-/-*^*/Pcgf2*^*fl/fl*^ ERT2-Cre ESCs ^59^ were treated with 800nM 4-hydroxytamoxifen (Sigma) for 96 hours or 72 hours, respectively. For RA differentiation of ESCs, 4×10^6^ ESCs were allowed to attach to gelatinised 15cm dishes for 6-8 hours and treated with 1μM all-trans retinoic acid (Sigma-Aldrich) in EC-10 medium (DMEM supplemented with 10% fetal bovine serum, L-Glutamine, beta-mercaptoethanol, non-essential amino acids and penicillin/streptomycin) for 48 hours. TOT2N E14 ESCs used for TetR targeting experiments were previously described ^69^.To generate TetR-CDK8 TOT2N ES line, TOT2N E14 ESCs were transfected using Lipofectamine 2000 (Thermo Fisher scientific) following manufacturer’s instructions. Stably-transfected cells were selected for 10 days using 1 μg/ml puromycin and individual clones were isolated and expanded in the presence of 1 μg/ml puromycin to maintain transgene expression. HEK293T cells, used for calibration of cross-linked cChIPseq experiments, were cultured in EC-10 media. All mammalian cell lines were cultured at 37°C and 5% CO2. SG4 Drosophila cells, used for calibration of ncRNAseq and native ChIPseq experiments, were grown at 25°C in Schneider’s medium (Thermo Fisher scientific) supplemented with 10% heat-inactivated fetal bovine serum (BioSera) and penicillin/streptomycin. All cell lines generated and grown in the Klose lab were routinely tested for mycoplasma infection.

### Generation of MED14-T7 *Med13/13l*^*fl/fl*^ ESC line

To allow for efficient chromatin immunoprecipitation of MED14, we introduced an N-terminal 3xT7-2xStrepII-FKBP12 tag to the endogenous *Med14* gene. The tag was synthesised by GeneArt (Thermo Fisher Scientific). The targeting construct was generated by Gibson assembly (Gibson Assembly Master Mix kit, New England Biolabs) of the PCR-amplified tag sequence and roughly 520bp homology arms surrounding the ATG start codon of the *Med14* gene, amplified from mouse genomic DNA.

The pSptCas9(BB)-2A-Puro(PX459)-V2.0 vector was obtained from Addgene (#62988) and the sgRNA was designed using the CRISPOR online tool (http://crispor.tefor.net/crispor.py). The targeting construct was designed such that the endogenous ATG sequence is removed and the Cas9 recognition site was disrupted by the insertion of the tag. ESCs were transfected in a single well of a 6-well plate with 0.5μg Cas9 guide plasmid and 2μg targeting construct plasmid using Lipofectamine 3000 (Thermo Fisher Scientific) according to manufacturer’s guidelines. The day after transfection, cells were passaged at a range of densities and subjected to puromycin selection (1 μg/ml) for 48 hours. Approximately 7-10 days following transfection, individual clones were isolated, expanded, and PCR-screened for the homozygous presence of the tag.

### Preparation of nuclear extracts and Western blot analysis

Harvested cells were resuspended in 10x pellet volume (PV) of Buffer A (10mM Hepes pH 7.9, 1.5mM MgCl2, 10mM KCl, 0.5mM DTT, 0.5mM PMSF, cOmplete protease inhibitor cocktail (Roche)) and incubated for 10 min at 4°C with slight agitation. After centrifugation, the cell pellet was resuspended in 3x PV Buffer A containing 0.1% NP-40 and incubated for 10 min at 4°C with slight agitation. Nuclei were recovered by centrifugation and the soluble nuclear fraction was extracted for 1 hr at 4°C with slight agitation using 1x PV Buffer C (10mM Hepes pH 7.9, 400mM NaCl, 1.5mM MgCl2, 26% glycerol, 0.2mM EDTA, cOmplete protease inhibitor cocktail). Protein concentration was measured using Bradford assay (BioRad).

Nuclear extract samples were resuspended with 1x SDS loading buffer (2% SDS, 0.1M Tris pH 6.8, 0.1M DTT, 10% glycerol, 0.1% bromophenolblue) and placed at 95°C for 5 mins. Between 25-35ug nuclear extract was separated on home-made SDS-PAGE gels or NuPAGE™ 3-8% Tris-Acetate gels (Life Technologies, for large Mediator subunits). Gels were blotted onto nitrocellulose membranes using the Trans-Blot Turbo transfer system (BioRad). Antibodies used for Western blot analysis were rabbit polyclonal anti-MED13L (A302-420A, Bethyl laboratories), rabbit polyclonal anti-MED13 (GTX129674, Genetex), rabbit monoclonal anti-CDK8 (ab229192, Abcam), rabbit polyclonal anti-CCNC (A301-989A, Bethyl laboratories), rabbit polyclonal anti-MED1 (A300-793A, Bethyl laboratories), rabbit polyclonal anti-MED15 (A302-422A, Bethyl laboratories), rabbit polyclonal anti-MED14 (A301-044A-T, Bethyl laboratories), rabbit monoclonal anti-RING1B (5694, Cell Signaling), rabbit monoclonal anti-SUZ12 (3737, Cell Signaling), rabbit polyclonal anti-PCGF2 (sc-10744, Santa Cruz), rabbit monoclonal anti-T7-Tag (D9E1X) (13246, Cell Signaling), mouse monoclonal anti-TBP (ab818, Abcam), rabbit monoclonal anti-HDAC1 (ab109411, Abcam), and mouse monoclonal anti-Flag (F1804, Sigma).

### Co-immunoprecipitation of the CDK-MED complex

For purification of the CDK-MED complex from wild type or tamoxifen-treated *Med13/13l*^*fl/fl*^ ESCs, 600μg of nuclear extract was diluted in BC150 buffer (50mM Hepes pH 7.9, 150 mM KCl, 0.5mM EDTA, 0.5mM DTT, cOmplete protease inhibitor cocktail (Roche)). Samples were incubated with 5μg CDK8 antibody (A302-500A, Bethyl laboratories) and 25U benzonase nuclease (Millipore) overnight at 4°C. Protein A agarose beads (RepliGen) were blocked for 1 hr at 4°C in Buffer BC150 containing 1% fish skin gelatine (Sigma) and 0.2 mg/ml BSA (New England Biolabs). The blocked beads were added to the samples and incubated for 4 hr at 4°C. Washes were performed using BC150 containing 0.02% NP-40. The beads were resuspended in 2x SDS loading buffer and boiled for 5 min to elute the immunoprecipitated complexes.

### Chromatin Immunoprecipitation

Chromatin immunoprecipitation was performed as described previously ^55^. In brief, 50×10^6^ ES cells were fixed for 45min with 2mM DSG (Thermo Fischer scientific) in PBS followed by 12.5 min with 1% formaldehyde (methanol-free, Thermo Fischer scientific). Reactions were quenched by the addition of glycine to a final concentration of 125 μM and the fixed cells were washed in ice-cold PBS and snap frozen in liquid nitrogen. 50×10^6^ HEK293T cells were fixed as above, snap frozen in 2×10^6^ aliquots, and stored at -80°C until further use.

For calibrated ChIPseq, 2×10^6^ HEK293T cells were resuspended in 1ml ice-cold lysis buffer (50mM HEPES pH 7.9, 140mM NaCl, 1 mM EDTA, 10% glycerol, 0.5% NP40, 0.25% Triton X-100) and added to 50×10^6^ fixed ESCs, resuspended in 9ml lysis buffer. The cell suspension was incubated for 10 min at 4°C. The released nuclei were then washed (10mM Tris-HCl pH 8, 200mM NaCl, 1mM EDTA, 0.5mM EGTA) for 10 min at 4°C. The chromatin pellet was resuspended in 1mL of ice-cold sonication buffer (10mM Tris-HCl pH 8, 100mM NaCl, 1mM EDTA, 0.5mM EGTA, 0.1% Na deoxycholate, 0.5% N-lauroylsarcosine) and sonicated for 25 cycles (30 sec on/off) using a BioRuptor Pico sonicator (Diagenode), shearing genomic DNA to produce fragments between 300bp and 1kb. Following sonication, Triton X-100 was added to a final concentration of 1%. Two hundred and fifty μg chromatin diluted 10-fold in ChIP dilution buffer (1% Triton-X100, 1mM EDTA, 20mM Tris-HCl (pH 8.0), 150mM NaCl) was used per IP. Three reactions were set up per condition to allow for maximal DNA recovery suitable for library preparation. Chromatin was precleared with protein A Dynabeads (Thermo Fischer scientific), blocked with 0.2 mg/ml BSA and 50 μg/ml yeast tRNA, and incubated with the respective antibodies overnight at 4°C. Antibody-bound chromatin was purified using blocked protein A Dynabeads for 3 hours at 4°C. ChIP washes were performed as described previously ^88^. ChIP DNA was eluted in ChIP elution buffer (1% SDS, 100mM NaHCO3) and reversed cross-linked overnight at 65°C with 200mM NaCl and RNase A (Sigma). The reverse cross-linked samples were treated with 20 μg/ml Proteinase K and purified using ChIP DNA Clean and Concentrator kit (Zymo research). The three reactions per condition were pooled at this stage. For each sample, corresponding Input DNA was also reverse cross-linked and purified. The efficiency of the ChIP experiments was confirmed by quantitative PCR. Prior to library preparation, 5-10ng ChIP material was diluted to 50ul in TLE buffer (10mM Tris-HCl pH8.0, 0.1mM EDTA) and sonicated with Bioruptor Pico sonicator for 17 min (30 sec on/off).

The antibodies used for ChIPseq experiments were rabbit polyclonal anti-CDK8 (A302-500A, Bethyl laboratories, 2.5μl), rabbit monoclonal anti-RING1B (5694, Cell Signaling, 3 μl), rabbit polyclonal anti-PCGF2 (sc-10744, Santa Cruz, 3ul), rabbit monoclonal anti-T7-Tag (D9E1X) (13246, Cell Signaling, 3ul). The antibodies used for ChIP-qPCR for TetO targeting experiments we rabbit polyclonal anti-FS2 (produced in house, ^88^, 3ul), polyclonal anti-MED12 (A300-774A, Bethyl laboratories, 3ul), polyclonal anti-MED1 (A300-793A, Bethyl laboratories, 3ul), polyclonal anti-CCNC (A301-989A, Bethyl laboratories, 3ul), and rabbit polyclonal anti-FS2 (produced in-house,, 5ul).

### Native Chromatin Immunoprecipitation

Native calibrated ChIPseq for H3K27me3 was performed as described previously ^59,88^. Briefly, 50×10^6^ ESCs were mixed with 20×10^6^ SG4 *Drosophila* cells and washed with 1x PBS prior to chromatin isolation. Nuclei were released in ice cold lysis buffer (10mM Tris-HCl pH 8.0, 10mM NaCl, 3mM MgCl2, 0.1% NP40), washed and resuspended in 1ml ice-cold digestion buffer (10mM Tris-HCl pH 8.0, 10mM NaCl, 3mM MgCl2, 0.1% NP40, 0.25M sucrose, 3mM CaCl2, 1x cOmplete protease inhibitor cocktail (Roche)). Chromatin was digested with 200U MNase (ThermoFischer scientific) for 5 min at 37°C and the reaction was stopped by the addition of 4mM EDTA pH 8.0. The samples were centrifuged at 1500g for 5 min at 4°C, the supernatant (S1) was retained. The remaining pellet was incubated with 300μl of nucleosome release buffer (10mM Tris-HCl pH 7.5, 10mM NaCl, 0.2mM EDTA, 1x protease inhibitor cocktail (Roche)) at 4°C for 1 h, passed five times through a 27G needle using a 1mL syringe, and spun at 1500g for 5 min at 4°C. The second supernatant (S2) was collected and combined with corresponding S1 sample from above. Digestion to mostly mononucleosomes was confirmed on a 1.5% agarose gel. The prepared native chromatin was aliquoted, snap frozen in liquid nitrogen and stored at -80°C until further use. ChIPs were performed as described previously ^59^, using 5ul of H3K27me3 antibody prepared in-house.

### Calibrated nuclear RNAseq

Nuclear RNA sample preparation was performed using 20×10^6^ ES or RA-treated cells and 8×10^6^ SG4 *Drosophila* cells as described previously ^59^. RNA was isolated from purified nuclei using RNeasy RNA extraction kit (Qiagen) and gDNA contamination was depleted using TURBO DNA-free Kit (ThermoFischer scientific). Quality of RNA was assessed using 2100 Bioanalyzer RNA 6000 Pico kit (Agilent).

### Library preparation and high-throughput sequencing

All cChIPseq experiments were performed in biological triplicates. All ncRNAseq experiments were performed in biological quadruplicates. Libraries for cChIPseq and Native cChIPseq were prepared from 5-10ng of ChIP and corresponding input DNA samples using NEBNext Ultra II DNA Library Prep Kit for Illumina (New England Biolabs), following manufacturer’s guidelines. For ncRNAseq, RNA samples (800ng) were depleted of rRNA using the NEBNext rRNA Depletion kit (New England Biolabs). RNA-seq libraries were prepared using the NEBNext Ultra Directional RNA Library Prep kit (New England Biolabs). Samples were indexed using NEBNext Multiplex Oligos (New England Biolabs). The average size and concentration of all libraries was analyzed using the 2100 Bioanalyzer High Sensitivity DNA Kit (Agilent) followed by qPCR quantification using SensiMix SYBR (Bioline, UK) and KAPA Illumina DNA standards (Roche). Libraries were sequenced as 40bp paired-end reads on Illumina NextSeq 500 platform.

### Massively parallel sequencing, data processing and normalization

For cChIP-seq, paired-end reads were aligned to concatenated mouse and spike-in genomes (mm10+hg19 for cross-linked cChIP-seq and mm10+dm6 for native cChIP-seq) using Bowtie 2 ^89^, with the ‘‘–no-mixed’’ and ‘‘–no-discordant’’ options specified. Reads that were mapped more than once were discarded, followed by removal of PCR duplicates using Sambamba ^90^.

For cnRNA-seq, paired-end reads were first aligned using Bowtie 2 (with ‘‘–very-fast,’’ ‘‘–no-mixed’’ and ‘‘–no-discordant’’ options) against the concatenated mm10 and dm6 rRNA genomic sequence (GenBank: BK000964.3 and M21017.1), to filter out reads mapping to rDNA fragments. All unmapped reads were then aligned against the genome sequence of concatenated mm10 and dm6 genomes using STAR ^91^. To improve mapping of intronic sequences of nascent transcripts abundant in nuclear RNA-seq, reads failing to map using STAR were aligned against the mm10+dm6 concatenated genome using Bowtie 2 (with’’–sensitive-local,’’ ‘‘–no-mixed’’ and ‘‘–no-discordant’’ options). PCR duplicates were removed using SAMTools ^92^.

For visualisation and annotation of genomic regions, internal normalisation of cChIPseq and ncRNAseq experiments was performed as described previously ^59^. In brief, mouse reads were randomly downsampled based on the spike-in ratio (hg19 or dm6) in each sample. To account for possible spike-in cell variation, the ratio of spike-in to mouse read counts in the corresponding ChIP inputs were used as a correction factors for cChIP-seq replicates. MED14-T7 ChIP-seq was performed without spike-in normalisation. Individual replicates were compared using multiBamSummary and plotCorrelation functions from deepTools (version 3.1.1) ^93^, confirming high degree of correlation (Pearson’s correlation coefficient > 0.9). Replicates were pooled for downstream analysis. Genome-coverage tracks for visualisation on the UCSC genome browser ^94^ were generated using the pileup function from MACS2 ^95^ for ChIP-seq and genomeCoverageBed from BEDtools (v2.17.0) ^96^ for cnRNA-seq.

### Read count quantitation and analysis

Heatmap and metaplot analysis for ChIP-seq was performed using computeMatrix and plotProfile/plotHeatmap functions from deepTools (v.3.1.1) ^93^, looking at read density at Polycomb domains, CDK8 peaks or TSS of CDK-MED-dependent genes. Intervals of interest were annotated with read counts from merged replicates, using a custom-made Perl script utilising SAMtools (v1.7) ^92^. Polycomb domains were defined in ^59^. CDK8 peaks were defined in ^55^. H3K27me3ac peaks were defined in ^41^.

For differential gene expression analysis, read counts were obtained from the non-normalised mm10 BAM files for a non-redundant mouse gene set, using a custom-made Perl script utilising SAMtools (v1.7) ^92^. The non-redundant mouse gene set (n=20,633) was obtained by filtering mm10 refGenes for very short genes with poor sequence mapability and highly similar transcripts. To identify significant changes in gene expression, a custom-made R script utilising DESeq2 ^97^ was used. For spike-in normalisation, read counts for the spike-in genome at a unique set of dm6 refGenes were supplied to calculate DESeq2 size factors which were then used for DESeq2 normalization of raw mm10 read counts, similarly to ^98^. For a change to be considered significant, a threshold of fold change > 1.5 and p-adj value < 0.05 was applied.

The distribution of log2-fold changes and normalised read counts at different genomics intervals was visualised using custom R scripts. For boxplot analyses, boxes show interquartile range (IQR) and whiskers extend by no more than 1.5xIQR.

### Hi-C library preparation and analysis

*In-situ* Hi-C in *Med13/13l*^*fl/fl*^ ESCs was performed in biological duplicates as described in ^99^. Hi-C libraries were sequenced on Illumina NextSeq 500 platform as 51bp or 40bp paired-reds. Hi-C sequencing data was mapped to GRCm38.p6 and processed with Hi-C-Pro 2.9 ^100.^Further data analysis was performed with GENOVA (github.com/deWitLab/GENOVA) ^101^.

TAD and loop coordinates of mouse ESC samples were taken from ^25^. Aggregate Peak and TAD analyses (APA; ATA) were performed on 10kb ice-normalized matrices with default parameters. PE-SCAn between the 100kb regions surrounding Ring1B peaks in Polycomb domains was also performed on these matrices. Super-enhancer coordinates for GRCm38.p6 were downloaded from dbSUPER ^102^. PE-SCAn between the 1Mb regions surrounding super-enhancers was performed using 20kb ice-normalized matrices, setting the bottom- and top-five percent values as outliers.

### Capture-C Extraction Protocol

Chromatin was extracted and fixed as described previously ^103^. Briefly, 10×10^6^ mouse ESCs were trypsinized, collected in 50ml falcon tubes in 9.3ml medium and crosslinked with 1.25ml 16% formaldehyde (1.89% final concentration; methanol-free, Thermo Fisher Scientific) while rotating for 10 min at room temperature. Cells were quenched with 1.5ml 1M cold glycine, washed with cold PBS and lysed for 20 minutes at 4°C in lysis buffer (10mM Tris pH 8, 10mM NaCl, 0.2% NP-40, supplemented with cOmplete proteinase inhibitors (Roche)) prior to snap freezing in 1ml lysis buffer on dry ice. Fixed chromatin was stored at -80°C.

### Capture-C Library Construction Protocol

Capture-C libraries were prepared as described previously ^104^. Briefly, lysates were thawed on ice, pelleted and resuspended in 650μl 1x DpnII buffer (New England Biology). Three 1.5ml tubes with 200μl lysate each were treated in parallel with 0.28% final concentration of SDS (Thermo Fischer Scientific) for 1hr at 37°C in a thermomixer shaking at 500rpm (30 sec on/off). Reactions were then quenched with 1.67% final concentration of Triton X-100 for 1hr at 37°C in a thermomixer shaking at 500rpm (30 sec on/off) and digested for 24 hours with 3×10μl DpnII (produced in-house) at 37°C in a thermomixer shaking at 500rpm (30 sec on/off). An aliquot from each reaction (100 μl) were taken for digestion control, reverse cross-linked and visualised on an agarose gel. The remaining chromatin was then independently ligated with 8μl T4 Ligase (240U, Themo Firsher Scientific) in a volume of 1440μl for 20 hours at 16°C. The nuclei containing ligated chromatin were pelleted to remove any non-nuclear chromatin, reverse cross-linked and the ligated DNA was phenol-chloroform purified. The sample was resupended in 300μl water and sonicated for 13 cycles (30sec on/off) using a Bioruptor Pico (Diagenode) to achieve a fragment size of approximately 200bp. Fragments were size-selected using AmpureX beads (Beckman Coulter), using selection ratios: 0.85x / 0.4x. Two reactions of 1-5μg DNA each were adaptor-ligated and indexed using NEBNext Ultra II DNA Library Prep Kit for Illumina (New England Biolabs) and NEBNext Multiplex Oligos for Illumina Primer sets 1 and 2 (New England Biolabs). The libraries were amplified with 7 PCR cycles using Herculase II Fusion Polymerase kit (Agilent). Libraries were next hybridized in the following way: For each promoter containing a DpnII restriction fragment, we designed two 70bp capture probes using the CapSequm online tool (http://apps.molbiol.ox.ac.uk/CaptureC/cgi-bin/CapSequm.cgi) with the following filtering parameters: Duplicates: <2, Density <30, SRepeatLength <30, Duplication: FALSE. For promoters for which no probes could be designed for the restriction fragment directly overlapping the TSS, probes were designed for the next-nearest DpnII fragment, if it was within 500bp of the TSS. The probes were pooled at 2.9nM each and the samples were multiplexed by mass prior to hybridization (2ug each, according to Qubit dsDNA BR Assay, Invitrogen). Hybridization was carried out using the Nimblegen SeqCap system (Roche, Nimblegen SeqCap EZ HE-oligo kit A #06777287001, Nimblegen SeqCap EZ HE-oligo kit B #06777317001, Nimblegen SeqCap EZ Accessory kit v2 #07145594001, Nimblegen SeqCap EZ Hybridisation and wash kit #05634261001) according to Roche protocol for 72 hours followed by a 24 hours hybridization (double Capture). Captured libraries were quantified by qPCR using SensiMix SYBR (Bioline, UK) and KAPA Illumina DNA standards (Roche) and sequenced on Illumina NextSeq 500 as 40bp paired-reads. Libraries for Capture-C in *Med13/13l*^*fl/f*^ and *Pcgf4*^-/-^*Pcgf2*^*fl/fl*^ were performed in biological triplicates (Capture set1) or biological duplicates (Capture set2, as control for captures in the TetR-fusion lines). Libraries for Capture-C in the TetR-fusion lines were performed in biological triplicates.

### Capture-C analysis

Paired-end reads were aligned to mm10 (or mm10 +BAC insert for TetR fusion cell lines) and filtered for Hi-C artefacts using HiCUP ^105^ and Bowtie 2 ^89^, with fragment filter set to 100-800bp. Read counts of reads aligning to captured gene promoters and interaction scores (=significant interactions) were then called by CHiCAGO ^106^.

For visualisation of Capture-C data, weighted pooled read counts from CHiCAGO data files were normalized to total read count aligning to captured gene promoters in the sample and further to the number of promoters in the respective capture experiment and multiplied by a constant number to simplify genome-browser visualization using the following formula: normCounts=1/cov*nprom*100000. Bigwig files were generated from these normalized read counts.

For comparative boxplot analysis we first determined all interactions between promoters and a given set of intervals (i.e. Polycomb domains) using Chicago score of >= 5 as a cutoff. Next, for each promoter-interval interaction we quantified the sum of normalized read counts or CHiCAGO scores across all DpnII fragments overlapping this interval. This number was then divided by the total number of interval-overlapping DpnII fragments to obtain mean normalized read counts/scores. For boxplot analyses, boxes show interquartile range (IQR) and whiskers show most extreme data point, which is no more than by 1.5xIQR.

## DATA AVAILABLITY

The datasets generated in this study are available from GEO database under accession number GSE185930.

## CODE AVAILABILITY

All R and Perl scripts used for data analysis in this study are available upon request. GENOVA is an open source software package available at http://www.github.com/deWitLab/GENOVA.

## ACKNOWLEDGEMENTS

We would like to thank Amy Hughes, Nadezda Fursova and Neil Blackledge for critical reading of the manuscript as well as members of the Klose lab for fruitful discussions. We thank Nadezda Fursova for help with DESeq2 analysis. We are grateful to Amanda Williams at the Department of Zoology, Oxford, for sequencing support on the NextSeq 500. Work in the Klose lab is supported by the Wellcome Trust (209400/Z/17/Z) and the European Research Council (681440). AF is supported by a Sir Henry Wellcome Post-doctoral fellowship. Work in the de Wit lab is supported by an ERC StG 637587 (‘HAP-PHEN’) and a Vidi grant from the Netherlands Scientific Organization (NWO, ‘016.16.316’).

## AUTHORS CONTRIBUTIONS

E.D. and R.J.K conceived the project and wrote the manuscript with contributions from all co-authors. E.D. performed experiments, data analysis and visualisation. A.F. and E.D. generated Capture-C libraries and A.F. performed Capture-C data analysis. K.D.F. generated Hi-C libraries and R.H.W. performed Hi-C data analysis. A.L. generated the MED14-T7 *Med13/13l*^*fl/fl*^ ESC line and libraries. E.W. and R.J.K supervised the project.

---

**Supplementary Figure 1:**
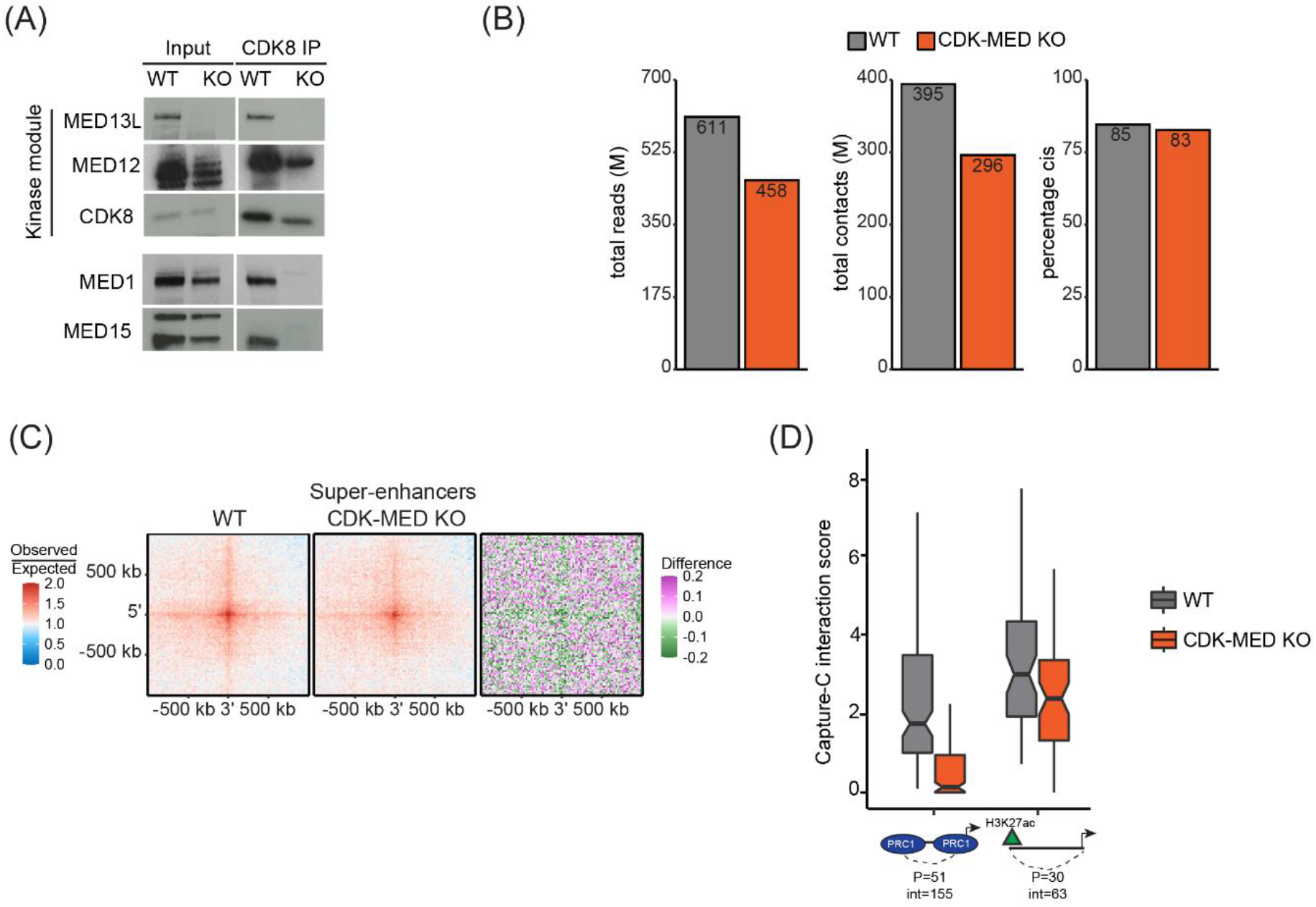
CDK-MED has a limited role in 3D genome organisation but is essential for Polycomb domain interactions. (A) Western blot analysis of CDK8 immunoprecipitation from nuclear extracts from *Med13/13l*^*fl/fl*^ (WT) and *Med13/13l*^*-/-*^ (CDK-MED KO) ESCs, probed with the indicated antibodies. (B) Quality control metrics of the Hi-C data, showing total sequenced read-pairs in millions, total valid contacts in millions and percentages *in cis* contacts for WT and CDK-MED KO ESCs. (C) Aggregate analysis of super enhancer interactions in WT and CDK-MED KO ESCs. The difference between WT and KO is shown. (D) Boxplot analysis of Capture-C interaction scores from WT and CDK-MED KO ESCs showing interactions between Polycomb gene promoters with other Polycomb-domains (left) or non-Polycomb gene promoters with active sites (H3K27ac, right). Number of promoters (P) and interactions (int) is shown.

**Supplementary Figure 2:**
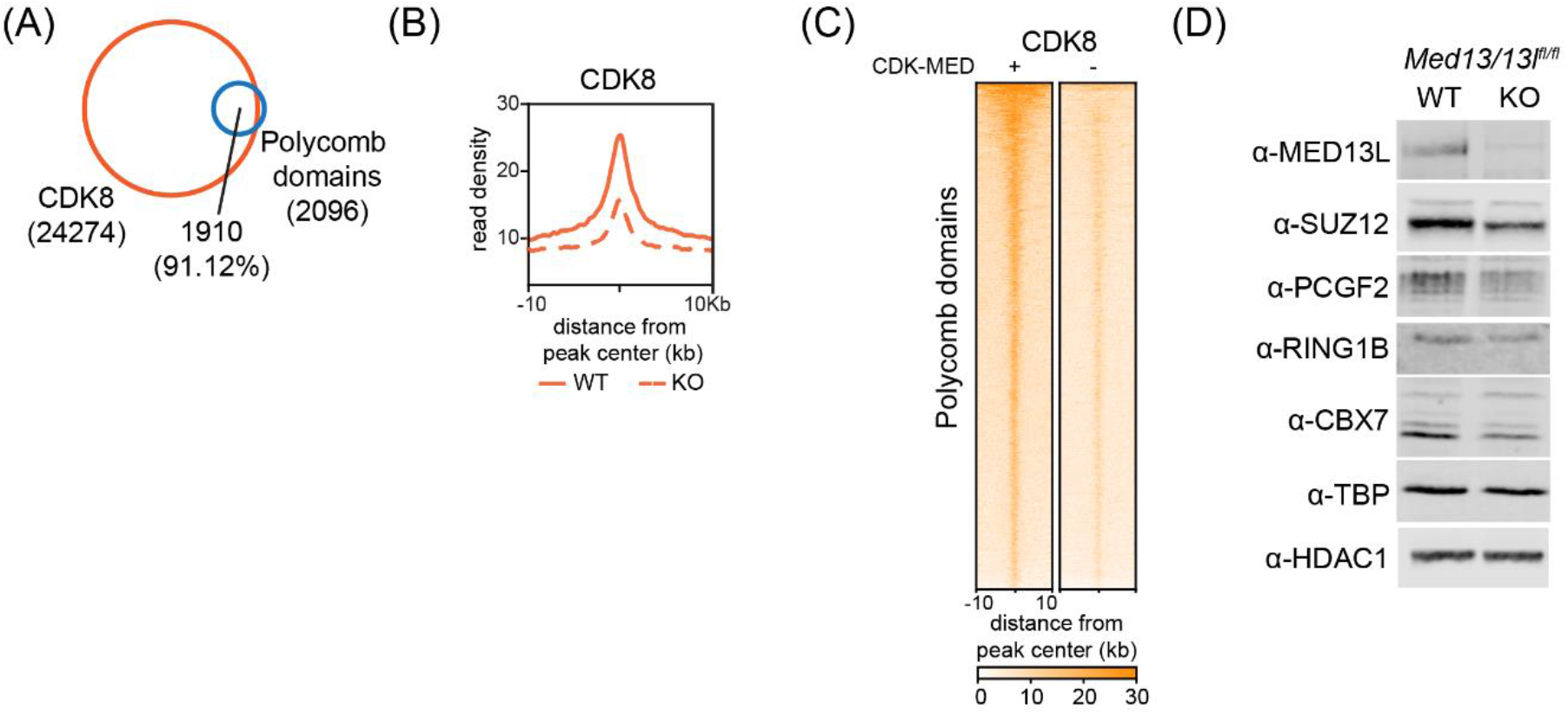
CDK-MED regulates canonical PRC1 binding. (A) A venn diagram showing the overlap between CDK8 peaks and Polycomb domains. Number of peaks and percent overlap are indicated. (B) Metaplot analysis of CDK8 enrichment at Polycomb domains (n=2097) in WT and CDK-MED KO ESCs. (C) Heatmaps showing CDK8 ChIPseq signal at Polycomb domains (n=2097) in WT and CDK-MED KO ESCs, sorted by decreasing RING1B signal. (D) Western blot analysis of nuclear extracts from WT and CDK-MED KO ESCs probed with the indicated antibodies. TBP and HDAC1 are used as loading controls.

**Supplementary Figure 3:**
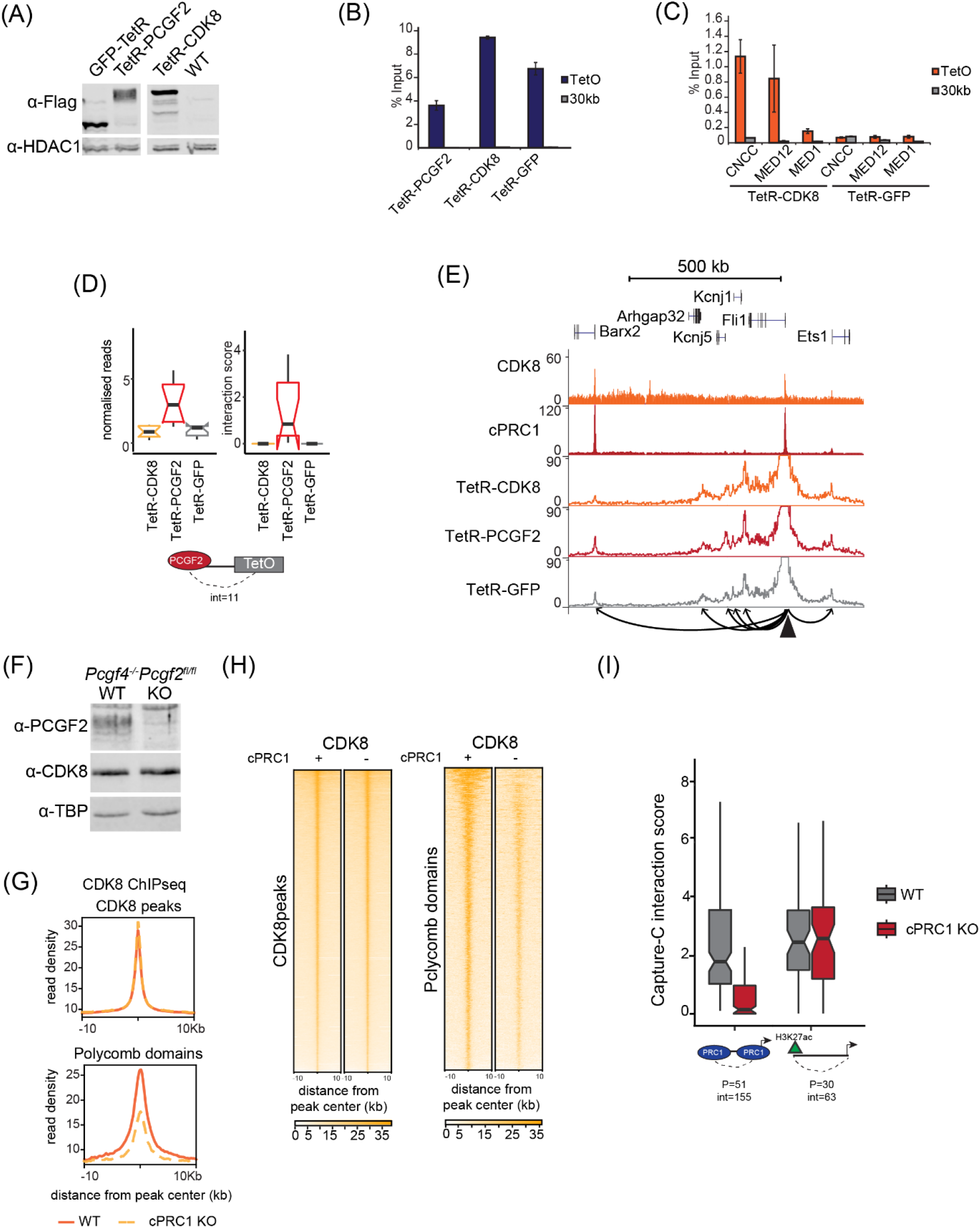
cPRC1 creates interactions between Polycomb domains. (A) Western blot analysis of nuclear extracts from the TetR-fusion lines used for Capture-C analysis probed with anti-Flag antibody to detect expression of the fusion proteins. HDAC1 is used as a loading control. (B) ChIP-qPCR analysis of binding of the different TetR-fusion lines to the TetO array. Error bars show standard deviation of three biological replicates. (C) ChIP-qPCR analysis of binding of the CDK-MED complex to the TetO array in the TetR-CDK8 and TetR-GFP lines. Error bars show standard deviation of three biological replicates. (D) Boxplot analysis of Capture-C mean normalised read counts and interaction scores in the TetR-fusion lines, looking at interactions with Polycomb domains (PCGF2-bound). Number of interactions is shown. (E) Snapshots showing Capture-C read count signal from TetR-CDK8, TetR-PCGF2 and TetR-GFP lines at a control locus. CDK8 and PCGF2 (cPRC1) ChIPseq signal is given as a reference. The *Fli1* promoter bait is shown as a triangle and interactions created with surrounding cPRC1-bound sites are represented with arrowheads. (F) Western blot analysis of nuclear extracts from WT and cPRC1 KO ESCs probed with the indicated antibodies. TBP is used as a loading control. (G) Metaplot analysis of CDK8 enrichment at CDK8 peaks (n=24275) and Polycomb domains (n=2097) in WT and cPRC1 KO ESCs. (H) Heatmaps showing CDK8 ChUPseq signal at CDK8 peaks (n=24275) and Polycomb domains (n=2097) in WT and cPRC1 KO ESCs, sorted by decreasing CDK8 or RING1B signal, respectively. (I) Boxplot analysis of Capture-C interaction scores from WT and cPRC1 KO ESCs showing interactions between Polycomb gene promoters with other Polycomb-domains (left) or non-Polycomb gene promoters with active sites (H3K27ac, right). Number of promoters (P) and interactions (int) is shown.

**Supplementary Figure 4:**
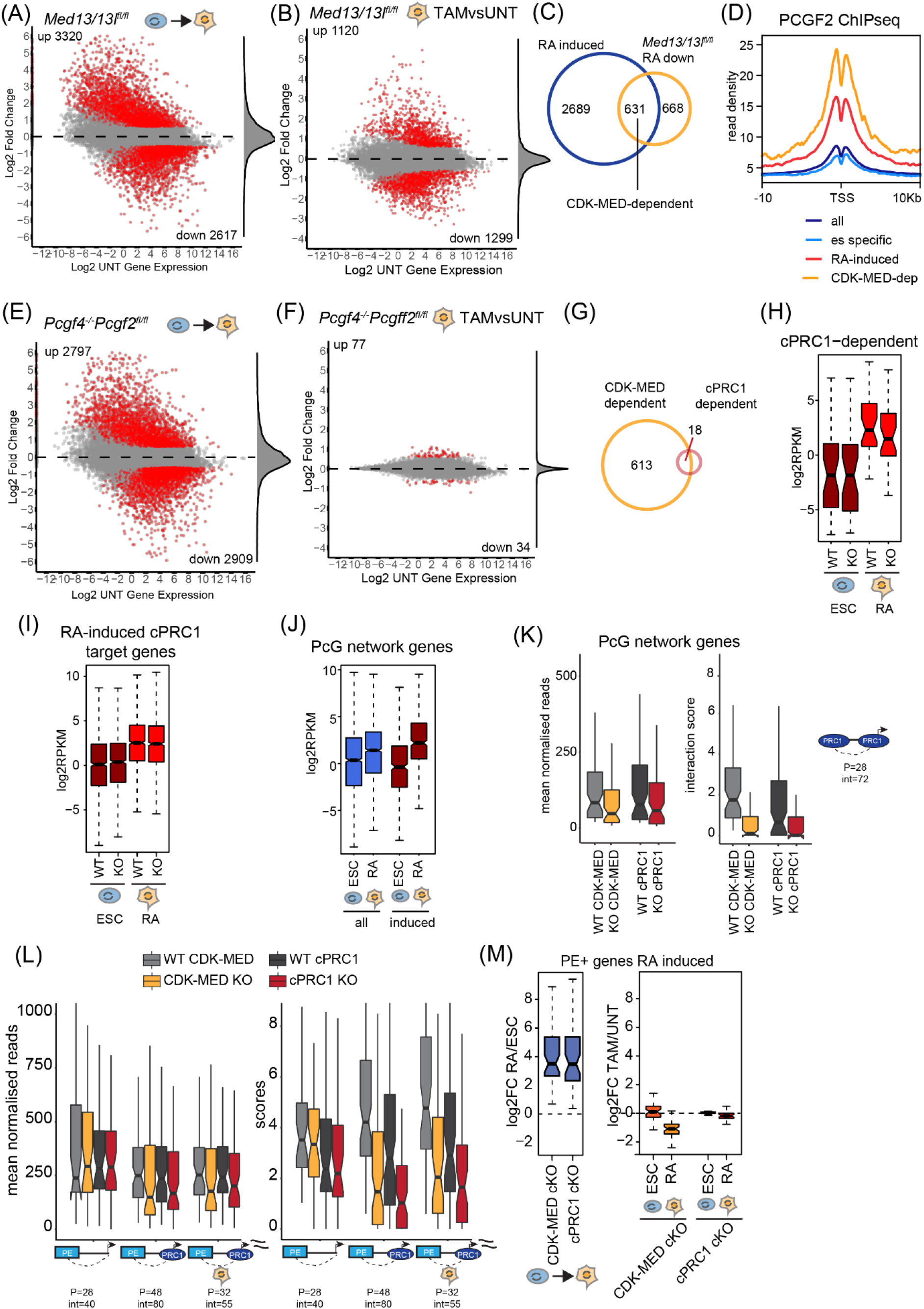
CDK-MED primes genes for activation during differentiation independently of cPRC1-mediated interactions. (A) An MA plot of log2 fold changes in gene expression (cnRNAseq) WT ESCs and RA-treated cells. Significant expression changes (>1.5 fold change and padj<0.05) are shown in red and number of genes is indicated. Distribution of gene expression changes is shown on the right as a density. (B) An MA plot of log2 fold changes in gene expression (cnRNAseq) WT and CDK-MED KO RA-treated cells. Significant expression changes (>1.5 fold change and padj<0.05) are shown in red and number of genes is indicated. Distribution of gene expression changes is shown on the right as a density. (C) A Venn diagram showing the overlap between RA-induced genes as defined in (A) and genes downregulated in CDK-MED KO cells, following RA treatment, as defined in (B). (D) A Metaplot showing enrichment of cPRC1 (PCGF2) over the transcription start site (TSS) of the indicated different classes of genes. All=20633, ES-specific=2617; RA-induced=3320; CDK-MED-dependent=631. (E) As in (A) for cPRC1 cKO cells. (F) As in (B) for cPRC1 cKO cells. (G) A Venn diagram showing the overlap between CDK-MED-dependent and cPRC1-dependent genes. Gene numbers are indicated. (H) Boxplot analysis of the expression of cPRC1-dependent genes (n=34), as defined in Figure S4F. (I) Boxplot analysis of the expression of RA-induced cPRC1 (PCGF2) target genes (n=1201) in WT and cPRC1 KO ESCs and following RA-induction. (J) Boxplot analysis of the expression of genes within the Polycomb network in ESCs and following RA induction (all=1974; RA-induced=482). (K) Boxplot analysis of Capture-C mean normalised read counts (left) and interaction score (right) from CDK-MED cKO and cPRC1 cKO ESCs showing interactions between promoters of genes within the Polycomb (PcG) network and Polycomb domains. Number of promoters (p) and interactions (int) is shown. (L) Boxplot analysis of Capture-C mean normalised read counts (left) and interaction score (right) from CDK-MED cKO and cPRC1 cKO ESCs showing interactions between gene promoters and poised enhancers (PE). Genes were divided into non-Polycomb targets (left set), Polycomb targets (middle set) and Polycomb targets induced by RA (right set). Number of promoters (P) and interactions (int) is shown. (M) Boxplot analysis of the expression of RA-induced genes that interact with a poised enhancer (n=55) in CDK-MED cKO and cPRC1 cKO cells. The difference between WT RA cells and ESCs (left), as well as KO and WT cells (right), is shown as log2FC.

**Supplementary Figure 5:**
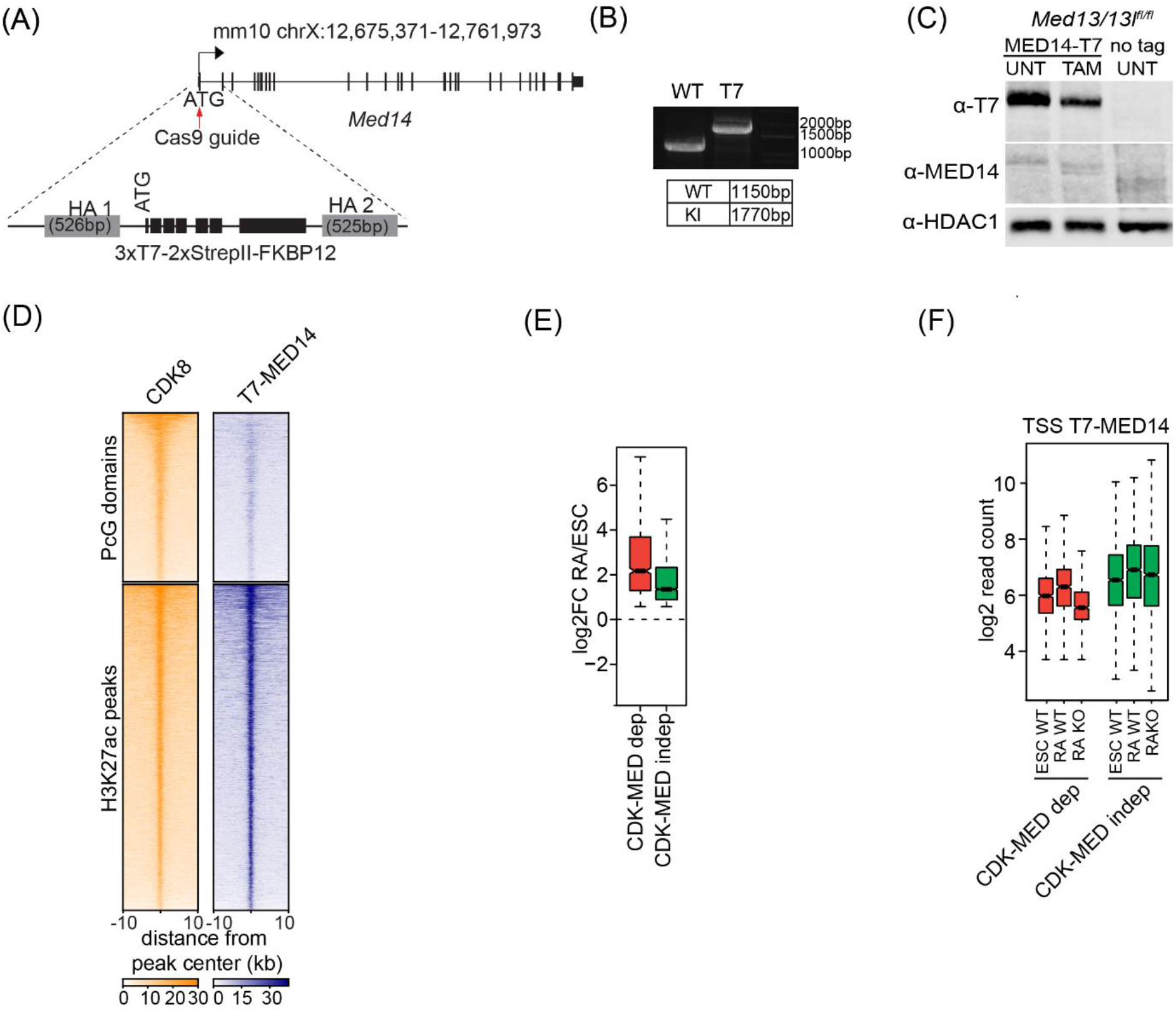
CDK-MED enables gene induction via recruitment of the Mediator complex. (A) A schematic illustration of the generation of the T7-MED14 expressing *Med13/13l*^*fl/fl*^ ESC line. (B) PCR showing amplification of homozygously-tagged T7-Med14 alleles. (C) Western blot analysis of nuclear extract from the T7-MED14 *Med13/13l*^*fl/fl*^ ESC line, following tamoxifen (TAM) treatment. Extract from an untagged ESC line was used as a control. HDAC1 was used as a loading control. (D) Heatmaps of CDK8 and T7-MED14 ChIPseq signal at Polycomb domains (n=2097) and H3K27ac peaks (n=4037), sorted by decreasing CDK8 signal. (E) Boxplots showing gene expression change (log2FC) of CDK-MED-dependent (n=631) and CDK-MED-independent (n=2689) RA-induced genes following RA differentiation of WT ESCs. (F) Boxplots showing T7-MED14 ChIPseq signal at the TSS (1000bp) of the different classes of RA-induced gene classes as defined in (E) in ESCs and RA-induced cells (WT and CDK-MED KO).

## Notes

### Competing Interest Statement

The authors have declared no competing interest.

## REFERENCES

1. Schoenfelder, S. & Fraser, P. Long-range enhancer-promoter contacts in gene expression control. Nat Rev Genet 20, 437–455 (2019).

2. de Laat, W. & Duboule, D. Topology of mammalian developmental enhancers and their regulatory landscapes. Nature 502, 499–506 (2013).

3. Furlong, E.E.M. & Levine, M. Developmental enhancers and chromosome topology. Science 361, 1341–1345 (2018).

4. Yokoshi, M. & Fukaya, T. Dynamics of transcriptional enhancers and chromosome topology in gene regulation. Dev Growth Differ 61, 343–352 (2019).

5. Rowley, M.J. & Corces, V.G. Organizational principles of 3D genome architecture. Nat Rev Genet 19, 789–800 (2018).

6. Rao, S.S.P. et al. Cohesin Loss Eliminates All Loop Domains. Cell 171, 305–320 e24 (2017).

7. Nora, E.P. et al. Targeted Degradation of CTCF Decouples Local Insulation of Chromosome Domains from Genomic Compartmentalization. Cell 169, 930–944 e22 (2017).

8. Hsieh, T.-H.S. et al. Enhancer-promoter interactions and transcription are maintained upon acute loss of CTCF, cohesin, WAPL, and YY1. 2021.07.14.452365 (2021).

9. Paliou, C. et al. Preformed chromatin topology assists transcriptional robustness of Shh during limb development. Proc Natl Acad Sci U S A 116, 12390–12399 (2019).

10. Liu, N.Q. et al. WAPL maintains a cohesin loading cycle to preserve cell-type-specific distal gene regulation. Nat Genet 53, 100–109 (2021).

11. Allen, B.L. & Taatjes, D.J. The Mediator complex: a central integrator of transcription. Nat Rev Mol Cell Biol 16, 155–66 (2015).

12. Jeronimo, C. et al. Tail and Kinase Modules Differently Regulate Core Mediator Recruitment and Function In Vivo. Mol Cell 64, 455–466 (2016).

13. Petrenko, N., Jin, Y., Wong, K.H. & Struhl, K. Mediator Undergoes a Compositional Change during Transcriptional Activation. Mol Cell 64, 443–454 (2016).

14. Jaeger, M.G. et al. Selective Mediator dependence of cell-type-specifying transcription. Nat Genet 52, 719–727 (2020).

15. Sun, F. et al. The Pol II preinitiation complex (PIC) influences Mediator binding but not promoter-enhancer looping. Genes Dev 35, 1175–1189 (2021).

16. Crump, N.T. et al. BET inhibition disrupts transcription but retains enhancer-promoter contact. Nat Commun 12, 223 (2021).

17. El Khattabi, L. et al. A Pliable Mediator Acts as a Functional Rather Than an Architectural Bridge between Promoters and Enhancers. Cell 178, 1145–1158 e20 (2019).

18. Blackledge, N.P. & Klose, R.J. The molecular principles of gene regulation by Polycomb repressive complexes. Nat Rev Mol Cell Biol (2021).

19. Schoenfelder, S. et al. Polycomb repressive complex PRC1 spatially constrains the mouse embryonic stem cell genome. Nat Genet 47, 1179–1186 (2015).

20. Kundu, S. et al. Polycomb Repressive Complex 1 Generates Discrete Compacted Domains that Change during Differentiation. Mol Cell 65, 432–446 e5 (2017).

21. Denholtz, M. et al. Long-range chromatin contacts in embryonic stem cells reveal a role for pluripotency factors and polycomb proteins in genome organization. Cell Stem Cell 13, 602–16 (2013).

22. Rhodes, J.D.P. et al. Cohesin Disrupts Polycomb-Dependent Chromosome Interactions in Embryonic Stem Cells. Cell Rep 30, 820–835 e10 (2020).

23. Vieux-Rochas, M., Fabre, P.J., Leleu, M., Duboule, D. & Noordermeer, D. Clustering of mammalian Hox genes with other H3K27me3 targets within an active nuclear domain. Proc Natl Acad Sci U S A 112, 4672–7 (2015).

24. Eskeland, R. et al. Ring1B compacts chromatin structure and represses gene expression independent of histone ubiquitination. Mol Cell 38, 452–64 (2010).

25. Bonev, B. et al. Multiscale 3D Genome Rewiring during Mouse Neural Development. Cell 171, 557–572 e24 (2017).

26. McLaughlin, K. et al. DNA Methylation Directs Polycomb-Dependent 3D Genome Reorganization in Naive Pluripotency. Cell Rep 29, 1974–1985 e6 (2019).

27. Bantignies, F. et al. Polycomb-dependent regulatory contacts between distant Hox loci in Drosophila. Cell 144, 214–26 (2011).

28. Joshi, O. et al. Dynamic Reorganization of Extremely Long-Range Promoter-Promoter Interactions between Two States of Pluripotency. Cell Stem Cell 17, 748–757 (2015).

29. Ogiyama, Y., Schuettengruber, B., Papadopoulos, G.L., Chang, J.M. & Cavalli, G. Polycomb-Dependent Chromatin Looping Contributes to Gene Silencing during Drosophila Development. Mol Cell 71, 73–88 e5 (2018).

30. Isono, K. et al. SAM domain polymerization links subnuclear clustering of PRC1 to gene silencing. Dev Cell 26, 565–77 (2013).

31. Moussa, H.F. et al. Canonical PRC1 controls sequence-independent propagation of Polycomb-mediated gene silencing. Nat Commun 10, 1931 (2019).

32. Cruz-Molina, S. et al. PRC2 Facilitates the Regulatory Topology Required for Poised Enhancer Function during Pluripotent Stem Cell Differentiation. Cell Stem Cell 20, 689–705 e9 (2017).

33. Crispatzu, G. et al. The chromatin, topological and regulatory properties of pluripotency-associated poised enhancers are conserved in vivo. Nat Commun 12, 4344 (2021).

34. Kondo, T. et al. Polycomb potentiates meis2 activation in midbrain by mediating interaction of the promoter with a tissue-specific enhancer. Dev Cell 28, 94–101 (2014).

35. Hug, C.B., Grimaldi, A.G., Kruse, K. & Vaquerizas, J.M. Chromatin Architecture Emerges during Zygotic Genome Activation Independent of Transcription. Cell 169, 216–228 e19 (2017).

36. Ing-Simmons, E. et al. Independence of chromatin conformation and gene regulation during Drosophila dorsoventral patterning. Nat Genet 53, 487–499 (2021).

37. Espinola, S.M. et al. Cis-regulatory chromatin loops arise before TADs and gene activation, and are independent of cell fate during early Drosophila development. Nat Genet 53, 477–486 (2021).

38. Ibrahim, D.M. & Mundlos, S. The role of 3D chromatin domains in gene regulation: a multi-facetted view on genome organization. Curr Opin Genet Dev 61, 1–8 (2020).

39. Benabdallah, N.S. et al. Decreased Enhancer-Promoter Proximity Accompanying Enhancer Activation. Mol Cell 76, 473–484 e7 (2019).

40. Alexander, J.M. et al. Live-cell imaging reveals enhancer-dependent Sox2 transcription in the absence of enhancer proximity. Elife 8(2019).

41. Feldmann, A., Dimitrova, E., Kenney, A., Lastuvkova, A. & Klose, R.J. CDK-Mediator and FBXL19 prime developmental genes for activation by promoting atypical regulatory interactions. Nucleic Acids Res 48, 2942–2955 (2020).

42. Kagey, M.H. et al. Mediator and cohesin connect gene expression and chromatin architecture. Nature 467, 430–5 (2010).

43. Phillips-Cremins, J.E. et al. Architectural protein subclasses shape 3D organization of genomes during lineage commitment. Cell 153, 1281–95 (2013).

44. Knuesel, M.T., Meyer, K.D., Bernecky, C. & Taatjes, D.J. The human CDK8 subcomplex is a molecular switch that controls Mediator coactivator function. Genes Dev 23, 439–51 (2009).

45. Knuesel, M.T., Meyer, K.D., Donner, A.J., Espinosa, J.M. & Taatjes, D.J. The human CDK8 subcomplex is a histone kinase that requires Med12 for activity and can function independently of mediator. Mol Cell Biol 29, 650–61 (2009).

46. Osman, S. et al. The Cdk8 kinase module regulates interaction of the mediator complex with RNA polymerase II. J Biol Chem 296, 100734 (2021).

47. Galbraith, M.D., Donner, A.J. & Espinosa, J.M. CDK8: a positive regulator of transcription. Transcription 1, 4–12 (2010).

48. Fant, C.B. & Taatjes, D.J. Regulatory functions of the Mediator kinases CDK8 and CDK19. Transcription 10, 76–90 (2019).

49. Donner, A.J., Szostek, S., Hoover, J.M. & Espinosa, J.M. CDK8 is a stimulus-specific positive coregulator of p53 target genes. Mol Cell 27, 121–33 (2007).

50. Donner, A.J., Ebmeier, C.C., Taatjes, D.J. & Espinosa, J.M. CDK8 is a positive regulator of transcriptional elongation within the serum response network. Nat Struct Mol Biol 17, 194–201 (2010).

51. D’Urso, A. et al. Set1/COMPASS and Mediator are repurposed to promote epigenetic transcriptional memory. Elife 5(2016).

52. Steinparzer, I. et al. Transcriptional Responses to IFN-gamma Require Mediator Kinase-Dependent Pause Release and Mechanistically Distinct CDK8 and CDK19 Functions. Mol Cell 76, 485–499 e8 (2019).

53. Chen, M. et al. CDK8/19 Mediator kinases potentiate induction of transcription by NFkappaB. Proc Natl Acad Sci U S A 114, 10208–10213 (2017).

54. Liu, Q. et al. The characterization of Mediator 12 and 13 as conditional positive gene regulators in Arabidopsis. Nat Commun 11, 2798 (2020).

55. Dimitrova, E. et al. FBXL19 recruits CDK-Mediator to CpG islands of developmental genes priming them for activation during lineage commitment. Elife 7(2018).

56. Papadopoulou, T., Kaymak, A., Sayols, S. & Richly, H. Dual role of Med12 in PRC1-dependent gene repression and ncRNA-mediated transcriptional activation. Cell Cycle 15, 1479–93 (2016).

57. Fukasawa, R., Iida, S., Tsutsui, T., Hirose, Y. & Ohkuma, Y. Mediator complex cooperatively regulates transcription of retinoic acid target genes with Polycomb Repressive Complex 2 during neuronal differentiation. J Biochem 158, 373–84 (2015).

58. Pavri, R. et al. PARP-1 determines specificity in a retinoid signaling pathway via direct modulation of mediator. Mol Cell 18, 83–96 (2005).

59. Fursova, N.A. et al. Synergy between Variant PRC1 Complexes Defines Polycomb-Mediated Gene Repression. Mol Cell 74, 1020–1036 e8 (2019).

60. Boyle, S. et al. A central role for canonical PRC1 in shaping the 3D nuclear landscape. Genes Dev 34, 931–949 (2020).

61. Wani, A.H. et al. Chromatin topology is coupled to Polycomb group protein subnuclear organization. Nat Commun 7, 10291 (2016).

62. Plys, A.J. et al. Phase separation of Polycomb-repressive complex 1 is governed by a charged disordered region of CBX2. Genes Dev 33, 799–813 (2019).

63. Grau, D.J. et al. Compaction of chromatin by diverse Polycomb group proteins requires localized regions of high charge. Genes Dev 25, 2210–21 (2011).

64. Lau, M.S. et al. Mutation of a nucleosome compaction region disrupts Polycomb-mediated axial patterning. Science 355, 1081–1084 (2017).

65. Francis, N.J., Kingston, R.E. & Woodcock, C.L. Chromatin compaction by a polycomb group protein complex. Science 306, 1574–7 (2004).

66. Cao, R. et al. Role of histone H3 lysine 27 methylation in Polycomb-group silencing. Science 298, 1039–43 (2002).

67. Min, J., Zhang, Y. & Xu, R.M. Structural basis for specific binding of Polycomb chromodomain to histone H3 methylated at Lys 27. Genes Dev 17, 1823–8 (2003).

68. Wang, L. et al. Hierarchical recruitment of polycomb group silencing complexes. Mol Cell 14, 637–46 (2004).

69. Blackledge, N.P. et al. Variant PRC1 complex-dependent H2A ubiquitylation drives PRC2 recruitment and polycomb domain formation. Cell 157, 1445–1459 (2014).

70. Wijchers, P.J. et al. Cause and Consequence of Tethering a SubTAD to Different Nuclear Compartments. Mol Cell 61, 461–473 (2016).

71. Malik, S. & Roeder, R.G. Dynamic regulation of pol II transcription by the mammalian Mediator complex. Trends Biochem Sci 30, 256–63 (2005).

72. Poss, Z.C., Ebmeier, C.C. & Taatjes, D.J. The Mediator complex and transcription regulation. Crit Rev Biochem Mol Biol 48, 575–608 (2013).

73. Soutourina, J. Transcription regulation by the Mediator complex. Nat Rev Mol Cell Biol 19, 262–274 (2018).

74. Rada-Iglesias, A. et al. A unique chromatin signature uncovers early developmental enhancers in humans. Nature 470, 279–83 (2011).

75. Ghavi-Helm, Y. et al. Enhancer loops appear stable during development and are associated with paused polymerase. Nature 512, 96–100 (2014).

76. Pachano, T. et al. Orphan CpG islands amplify poised enhancer regulatory activity and determine target gene responsiveness. Nat Genet 53, 1036–1049 (2021).

77. Loubiere, V., Papadopoulos, G.L., Szabo, Q., Martinez, A.M. & Cavalli, G. Widespread activation of developmental gene expression characterized by PRC1-dependent chromatin looping. Sci Adv 6, eaax4001 (2020).

78. Sugishita, H. et al. Variant PCGF1-PRC1 links PRC2 recruitment with differentiation-associated transcriptional inactivation at target genes. Nat Commun 12, 5341 (2021).

79. Akasaka, T. et al. Mice doubly deficient for the Polycomb Group genes Mel18 and Bmi1 reveal synergy and requirement for maintenance but not initiation of Hox gene expression. Development 128, 1587–97 (2001).

80. Isono, K. et al. Mammalian polyhomeotic homologues Phc2 and Phc1 act in synergy to mediate polycomb repression of Hox genes. Mol Cell Biol 25, 6694–706 (2005).

81. Miao, Y.L. et al. Mediator complex component MED13 regulates zygotic genome activation and is required for postimplantation development in the mouse. Biol Reprod 98, 449–464 (2018).

82. Westerling, T., Kuuluvainen, E. & Makela, T.P. Cdk8 is essential for preimplantation mouse development. Mol Cell Biol 27, 6177–82 (2007).

83. Rocha, P.P., Scholze, M., Bleiss, W. & Schrewe, H. Med12 is essential for early mouse development and for canonical Wnt and Wnt/PCP signaling. Development 137, 2723–31 (2010).

84. Gaytan de Ayala Alonso, A. et al. A genetic screen identifies novel polycomb group genes in Drosophila. Genetics 176, 2099–108 (2007).

85. Postlmayr, A., Dumeau, C.E. & Wutz, A. Cdk8 is required for establishment of H3K27me3 and gene repression by Xist and mouse development. Development 147(2020).

86. McCleland, M.L. et al. Cdk8 deletion in the Apc(Min) murine tumour model represses EZH2 activity and accelerates tumourigenesis. J Pathol 237, 508–19 (2015).

87. Scelfo, A. et al. Functional Landscape of PCGF Proteins Reveals Both RING1A/B-Dependent- and RING1A/B-Independent-Specific Activities. Mol Cell 74, 1037–1052 e7 (2019).

88. Farcas, A.M. et al. KDM2B links the Polycomb Repressive Complex 1 (PRC1) to recognition of CpG islands. Elife 1, e00205 (2012).

89. Langmead, B. & Salzberg, S.L. Fast gapped-read alignment with Bowtie 2. Nat Methods 9, 357–9 (2012).

90. Tarasov, A., Vilella, A.J., Cuppen, E., Nijman, I.J. & Prins, P. Sambamba: fast processing of NGS alignment formats. Bioinformatics 31, 2032–4 (2015).

91. Dobin, A. et al. STAR: ultrafast universal RNA-seq aligner. Bioinformatics 29, 15–21 (2013).

92. Li, H. et al. The Sequence Alignment/Map format and SAMtools. Bioinformatics 25, 2078–9 (2009).

93. Ramirez, F. et al. deepTools2: a next generation web server for deep-sequencing data analysis. Nucleic Acids Res 44, W160–5 (2016).

94. Kent, W.J. et al. The human genome browser at UCSC. Genome Res 12, 996–1006 (2002).

95. Zhang, Y. et al. Model-based analysis of ChIP-Seq (MACS). Genome Biol 9, R137 (2008).

96. Quinlan, A.R. BEDTools: The Swiss-Army Tool for Genome Feature Analysis. Curr Protoc Bioinformatics 47, 11 12 1–34 (2014).

97. Love, M.I., Huber, W. & Anders, S. Moderated estimation of fold change and dispersion for RNA-seq data with DESeq2. Genome Biol 15, 550 (2014).

98. Taruttis, F. et al. External calibration with Drosophila whole-cell spike-ins delivers absolute mRNA fold changes from human RNA-Seq and qPCR data. Biotechniques 62, 53–61 (2017).

99. Haarhuis, J.H.I. et al. The Cohesin Release Factor WAPL Restricts Chromatin Loop Extension. Cell 169, 693–707 e14 (2017).

100. Servant, N. et al. HiC-Pro: an optimized and flexible pipeline for Hi-C data processing. Genome Biol 16, 259 (2015).

101. van der Weide, R.H. et al. Hi-C analyses with GENOVA: a case study with cohesin variants. NAR Genom Bioinform 3, lqab040 (2021).

102. Khan, A. & Zhang, X. dbSUPER: a database of super-enhancers in mouse and human genome. Nucleic Acids Res 44, D164–71 (2016).

103. Hughes, J.R. et al. Analysis of hundreds of cis-regulatory landscapes at high resolution in a single, high-throughput experiment. Nat Genet 46, 205–12 (2014).

104. Downes, D.J. et al. High-resolution targeted 3C interrogation of cis-regulatory element organization at genome-wide scale. Nat Commun 12, 531 (2021).

105. Wingett, S. et al. HiCUP: pipeline for mapping and processing Hi-C data. F1000Res 4, 1310 (2015).

106. Cairns, J. et al. CHiCAGO: robust detection of DNA looping interactions in Capture Hi-C data. Genome Biol 17, 127 (2016).

